# The genomic and transcriptional landscape of the spinal cord H3K27-altered diffuse midline glioma suggests the potential therapeutic strategy

**DOI:** 10.1101/2025.05.25.655886

**Authors:** Yiwei Xiao, Mengyao Li, Qiang Gao, Zhihong Qian, Yukui Shang, Linkai Jing, Zili Zhen, Yong Ai, Guihuai Wang, Kehkooi Kee, Wei Zhang

## Abstract

Diffuse midline glioma with H3K27-altered (DMG) is a lethal pediatric malignancy that primarily occurs in the brainstem of children. In adults, DMG arising in the spinal cord is rarely reported and remains poorly understood. The cellular architecture and clinical outcomes of DMG vary depending on the tumor’s location. To dissect the genomic and transcriptional architecture of a cohort of spinal cord DMG patients, we conducted whole-exome sequencing and single-cell RNA sequencing on 19 tumor samples from 7 patients. We identified previously underappreciated stem-like cells associated with hypoxia in malignant tumor cells, which were present in all samples of spinal cord DMG but absent in published intracranial DMG, corroborating cellular heterogeneity based on tumor location. We discovered that tumor-associated myeloid cells constituted the largest population of non-malignant cells in adult spinal cord DMG. Our research revealed a transition from microglia to macrophages, characterized by enhanced cell-to-cell interactions mediated through the VEGF signaling pathway network. Notably, we found that epigenetic agents can repress the expression of hypoxia-related gene programs in patient-derived spinal cord DMG cultures and inhibit cell proliferation in vitro. Collectively, our study reveals a crucial anatomical dimension that DMG exhibits a location-specific genomic and transcriptional landscape despite shared H3K27 alterations, providing the paradigm for developing precision therapeutic strategies that extend beyond DMG to other molecularly defined yet anatomically divergent malignancies.

## Introduction

Diffuse midline glioma with H3K27-altered (DMG) is an epigenomic reprogramming-driven malignancy in the central nervous system [1, 2]. Typically, a lysine-to-methionine substitution at position 27 of the histone H3 (H3K27M) occurs in either non-canonical (H3.3) in over 70% of cases or canonical (H3.1 or H3.2) histone H3 isoforms. The presence of H3K27M inhibits EZH2 (Ehhancer of zeste homolog 2), a subunit of PRC2 (Polycomb repressive complex 2) that is responsible for establishing H3K27 methylation. This inhibition results in a widespread loss of H3K27me3 methylation [1, 3, 4]. These post-translational modifications (PTMs) have a dominant effect, triggering tumorigenesis by enhancing self-renewal, disrupting differentiation, and initiating tumor formation [1, 5, 6]. Hitherto, established treatment guidelines remain absent and clinical trials are preferred. In 2023, we presented our initial application of epigenetic agent-based immunotherapy in treating four adult recurrent and progressive DMG patients, including two patients with spinal cord DMG and two with thalamic DMG [7]. Remarkably, one spinal cord DMG patient with leptomeningeal metastasis (LM) survived for 20 months from diagnosis and 16 months following the initiation of epigenetic agent-based immunotherapy with improved quality of life. In contrast, the median overall survival for high-grade glioma patients with LM is only 1.6-3.8 months, as reported in the literature [7].

Historically, the understanding of DMG has primarily come from studies on pediatric pontine tumors due to their higher prevalence [8–10]. However, in recent years, there has been growing recognition of adult DMG in other midline structures. The prognosis of DMG is generally poor and varies depending on the subtype of histone H3 mutation and the specific midline anatomical locations involved. Patients harboring H3.3K27M mutation have a significantly shorter overall survival of merely 11 months, compared to those bearing H3.1/3.2K27M mutation who have an overall survival of 16 months [2, 11–13]. Meanwhile, patients with pontine DMG generally have the worst overall survival of only 6-9 months compared to those with DMG located in the thalamus or spinal cord [14, 15] This disparity indicates that different mutation subtypes and anatomical locations contribute to the heterogeneity of DMG, leading to varied prognosis for patients in clinical settings. Moreover, single-cell RNA sequencing (scRNA-seq) analysis, primarily focused on pediatric pontine DMG, has revealed that the origin of these cells could be neural stem cells that have adopted a transcriptomic characteristic of oligodendrocyte precursor cells (OPC). These OPC-like cells are partially sustained by *PDGFRA* signaling, which plays a crucial role in gliomagenesis and may serve as a potential therapeutic target [16]. Additionally, a comprehensive study that conducted multi-omic profiling of 50 DMG tumors, predominantly intracranial tumors along with 2 pediatric spinal cord DMG, implied that OPC-like cells are stem-like cells present across all midline anatomical locations, albeit with varying levels of maturation [17]. Together, OPC-like cells seem to play a pivotal role in oncohistone-driven tumorigenesis and could serve as potential therapeutic targets in the treatment of DMG.

However, both we and others reported that DMG occurs even more rarely in the spinal cord, primarily affecting adults [7, 10, 12, 18, 19]. In the presence of a shared H3K27M mutation, spinal cord DMG exhibits both intrinsic and extrinsic heterogeneity due to its location within the narrow and elongated osseous spinal canal architecture [12, 20–22]. For decades, spinal cord DMG has been significantly underappreciated and overlooked as a distinct subject of study, especially compared to its intracranial counterparts [16, 17]. Consequently, delineating spinal cord DMG across multiple tumor sites and charting them on single-cell and molecular levels will help bridge the gap in understanding this rare condition and may shed light on developing precise therapeutic strategies according to histone mutation-based, anatomically-specific DMG in clinical practice.

Here, we dissected the genomic and transcriptional signatures of spinal cord DMG by integrating whole-exome sequencing (WES) and scRNA-seq analyses of 19 tumor specimens from 7 patients (Figure 1). We identified a subset of malignant cells defined by differential expression of a hypoxia-related gene program, which was absent in intracranial H3K27-altered DMG [16]. These cells displayed a stem-like phenotype similar to OPC-like cells that were concurrently present in spinal DMG. Meanwhile, we observed a state transition of tumor-associated microglia to macrophages, accompanied by increased expression of hypoxia-related genes and activation of functional pathways within a tumor microenvironment characterized by extensive cell-to-cell interactions via the VEGF signaling pathway. Importantly, the combined application of epigenetic agents, as reported in our previous study [7], led to a downregulated hypoxia-related gene program and suppressed the proliferation of patient-derived spinal cord DMG cells *in vitro*. This suggests a potential therapeutic strategy for treating these tumors.

**Figure 1.**
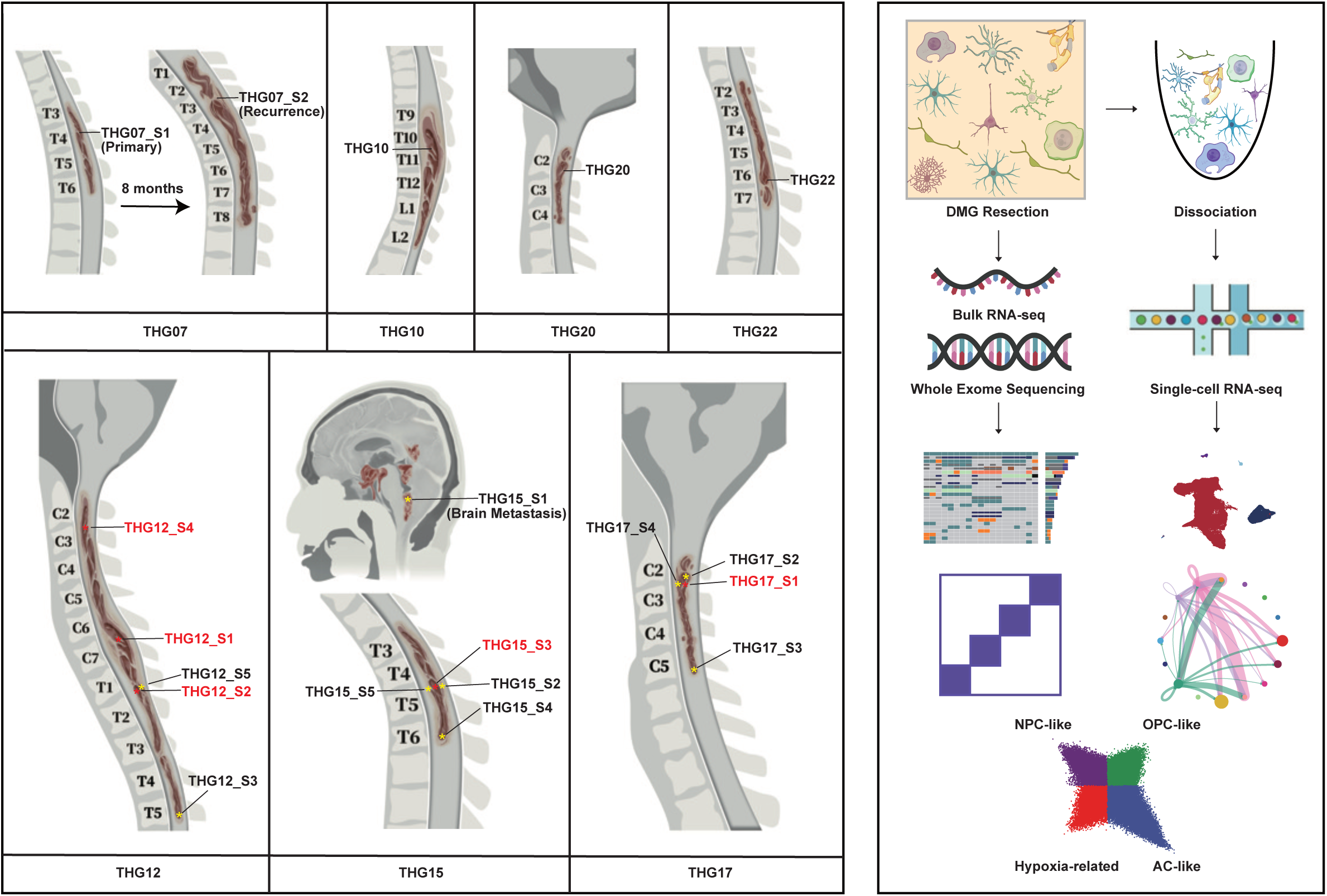
Schematic of overall workflow. The location of each tumor sample is indicated in the left panel. When multiple samples from the same lesion are present, core tumor samples are highlighted in red. Details of clinical characteristics of the patients are listed in Table 1. Tumor samples were subjected to analyses either as bulk tissues or single-cell suspensions as indicated in the right panel.

**Table 1.**
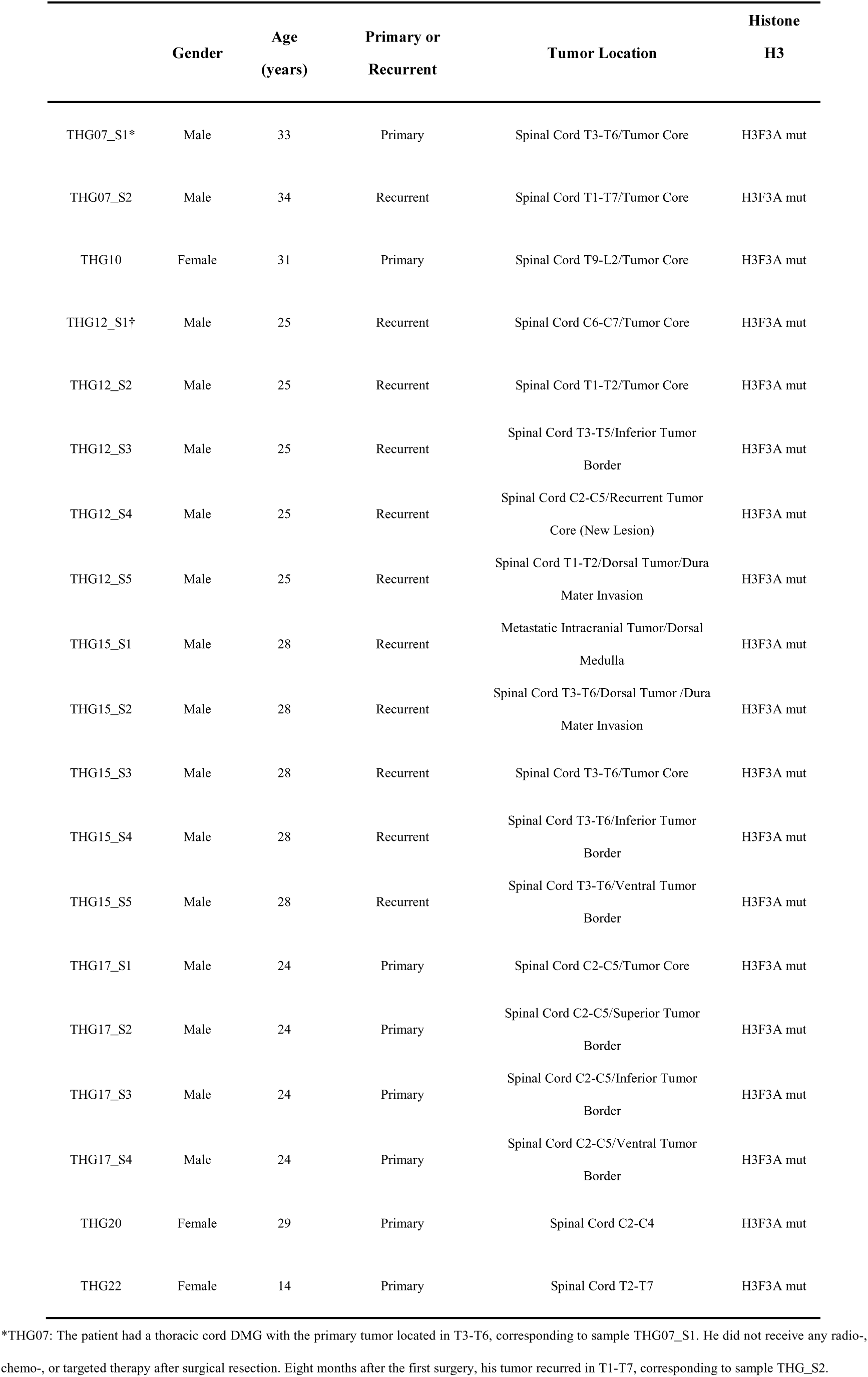
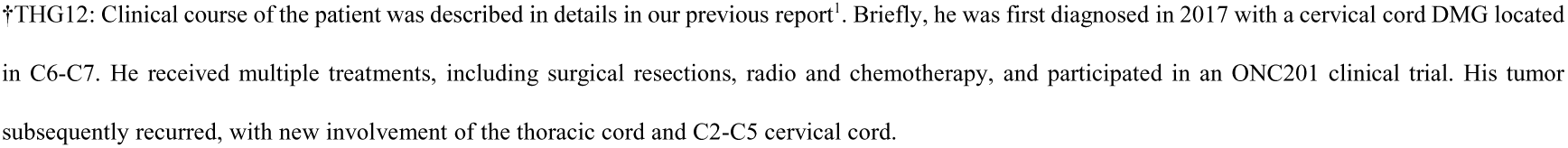
Summary of Clinical Parameters of Patients.

|  | Gender | Age<br>(years) | Primary or<br>Recurrent | Tumor Location | Histone<br>H3 |
| --- | --- | --- | --- | --- | --- |
| THG07_S1* | Male | 33 | Primary | Spinal Cord T3-T6/Tumor Core | H3F3A mut |
| THG07_S2 | Male | 34 | Recurrent | Spinal Cord T1-T7/Tumor Core | H3F3A mut |
| THG10 | Female | 31 | Primary | Spinal Cord T9-L2/Tumor Core | H3F3A mut |
| THG12_S1† | Male | 25 | Recurrent | Spinal Cord C6-C7/Tumor Core | H3F3A mut |
| THG12_S2 | Male | 25 | Recurrent | Spinal Cord T1-T2/Tumor Core | H3F3A mut |
| THG12_S3 | Male | 25 | Recurrent | Spinal Cord T3-T5/Inferior Tumor<br>Border | H3F3A mut |
| THG12_S4 | Male | 25 | Recurrent | Spinal Cord C2-C5/Recurrent Tumor<br>Core (New Lesion) | H3F3A mut |
| THG12_S5 | Male | 25 | Recurrent | Spinal Cord T1-T2/Dorsal Tumor/Dura<br>Mater Invasion | H3F3A mut |
| THG15_S1 | Male | 28 | Recurrent | Metastatic Intracranial Tumor/Dorsal<br>Medulla | H3F3A mut |
| THG15_S2 | Male | 28 | Recurrent | Spinal Cord T3-T6/Dorsal Tumor /Dura<br>Mater Invasion | H3F3A mut |
| THG15_S3 | Male | 28 | Recurrent | Spinal Cord T3-T6/Tumor Core | H3F3A mut |
| THG15_S4 | Male | 28 | Recurrent | Spinal Cord T3-T6/Inferior Tumor<br>Border | H3F3A mut |
| THG15_S5 | Male | 28 | Recurrent | Spinal Cord T3-T6/Ventral Tumor<br>Border | H3F3A mut |
| THG17_S1 | Male | 24 | Primary | Spinal Cord C2-C5/Tumor Core | H3F3A mut |
| THG17_S2 | Male | 24 | Primary | Spinal Cord C2-C5/Superior Tumor<br>Border | H3F3A mut |
| THG17_S3 | Male | 24 | Primary | Spinal Cord C2-C5/Inferior Tumor<br>Border | H3F3A mut |
| THG17_S4 | Male | 24 | Primary | Spinal Cord C2-C5/Ventral Tumor<br>Border | H3F3A mut |
| THG20 | Female | 29 | Primary | Spinal Cord C2-C4 | H3F3A mut |
| THG22 | Female | 14 | Primary | Spinal Cord T2-T7 | H3F3A mut |

\*THG07: The patient had a thoracic cord DMG with the primary tumor located in T3-T6, corresponding to sample THG07\_S1. He did not receive any radio-, chemo-, or targeted therapy after surgical resection. Eight months after the first surgery, his tumor recurred in T1-T7, corresponding to sample THG\_S2.
†THG12: Clinical course of the patient was described in details in our previous report<sup>1</sup>. Briefly, he was first diagnosed in 2017 with a cervical cord DMG located in C6-C7. He received multiple treatments, including surgical resections, radio and chemotherapy, and participated in an ONC201 clinical trial. His tumor subsequently recurred, with new involvement of the thoracic cord and C2-C5 cervical cord.

## Results

### The survival in patients with spinal cord H3K27-altered DMG compared with intracranial tumors

The survival of spinal cord H3K27-altered DMG is less well characterized than that of their intracranial counterparts due to their rarity [10, 23]. To evaluate the prognosis of our patient population, we gathered clinical data from 38 cases of spinal cord H3K27-altered DMG that were surgically treated or biopsied at our hospital from 2015 to 2021, and assessed the overall survival using Kaplan-Meier analysis (Figure 2; Supplementary information, Table S1). This cohort included 9 pediatric (<18 years of age) and 29 adult patients, resulting in a male-to-female ratio of 1.53 (23 males and 15 females). The overall survival of these patients ranged from 1 to 87 months, with a median survival of 17.0 months, which is consistent with previous reports [24–26]. To compare the survival outcomes between spinal and intracranial H3K27-altered DMG, we compiled data from the literature, including 79 cases of brainstem tumors and 31 cases in the thalamus. These patients received either conventional treatments, such as surgical resection, radiotherapy, temozolomide, and/or bevacizumab or were treatment-naïve (i.e., underwent biopsy only) [6, 22, 27–29] (Supplementary information, Table S1). The median overall survival for brainstem and thalamic H3K27-altered DMG was 8.5 and 10.4 months, respectively. Both of these are significantly shorter than the median survival for spinal cord DMG (17.0 vs. 8.5 months, hazard ratio, 0.49, 95% confidence interval, 0.32-0.75, P =0.0003; 17.0 vs. 10.4 months, hazard ratio, 0.5668, 95% confidence interval, 0.31-1.04, P =0.0294) (Figure 2). Together, these findings support the survival variability of H3K27-altered DMG based on tumor location, with the longest survival observed in patients with spinal cord DMG.

**Figure 2.**
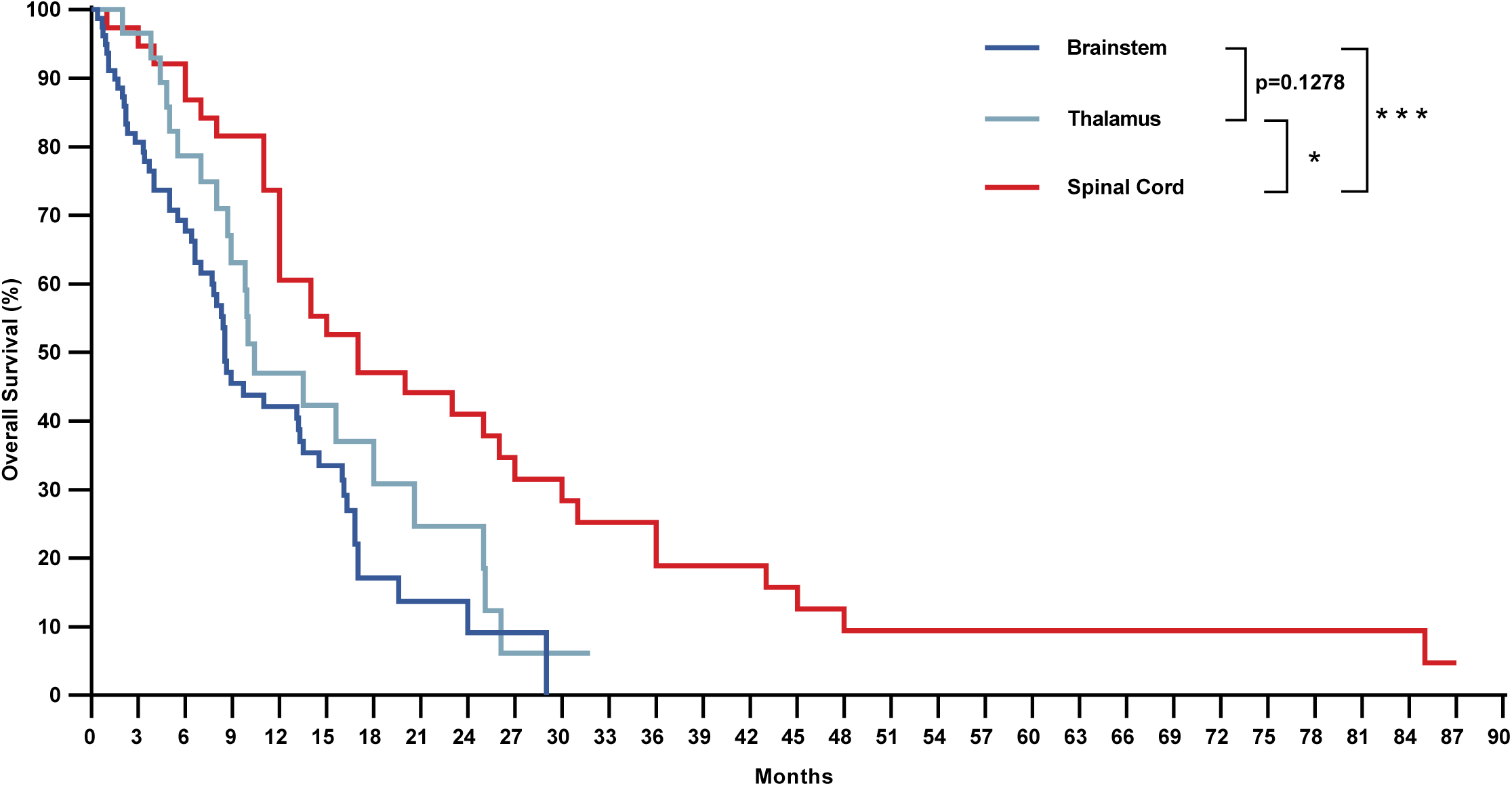
Kaplan-Meier analysis of patient survival with H3K27-altered DMG in different locations. Overall survival of patients with spinal cord H3K27-altered DMG treated at our hospital from 2015 to 2021 was analyzed and compared with that of patients with brainstem and thalamus DMG from literature data. *, p<0.05, ***, p<0.001.

### Hypoxia-related gene programs were extensively and specifically expressed in spinal cord H3K27-altered DMG tumor cells

Given the location-based variation in patient survival, we speculated that the genomic landscape of spinal cord H3K27-altered DMG might differ from that of the intracranial counterparts. We examined 19 tumor samples from 7 patients with spinal cord H3K27-altered DMG using WES and scRNA-seq analysis (Figure 1; Detailed clinical information was provided in Table 1). The genomic landscapes of all samples were summarized in Figure 3A. Notably, the K27M mutation occurred in H3F3A in all 19 samples. We identified multiple concurrent oncogenic mutations commonly associated with H3K27-altered DMG [30], such as mutations in TP53 (13 out of 19 samples from 5 out of 7 patients), PDGFRA (5 out of 19 samples from 1 out of 7 patients), PPM1D (11 out of 19 samples from 3 out of 7 patients), and FGFR1 (5 out of 19 samples from 1 out of 7 patients), with considerable consistency among tumors originating from the same patient (Figure 3A, Table S2). We utilized GISTIC (genomic identification of significant targets in cancer) to determine copy-number drivers based on focality, amplitude, and recurrence of alterations (CNA). Multiple well-known cancer driver genes were targeted by those focal CNAs, including amplifications in PDGFRA (58%) and CDK4 (47%), as well as deletions in PTEN (47%) and CDKN2A/B (26%) (Figure 3A). Among these, the patient THG15 harbored a DNA loss in the CDKN2A/B locus. CDKN2A encodes the tumor suppressor p16, which plays a critical role in regulating the cell cycle via inhibiting the cyclin-dependent kinase 4/6 (CDK4/6) [31]. We observed frequent deletions in PTEN, which were rarely detected in other previously reported DMG patients[17, 32].

**Figure 3.**
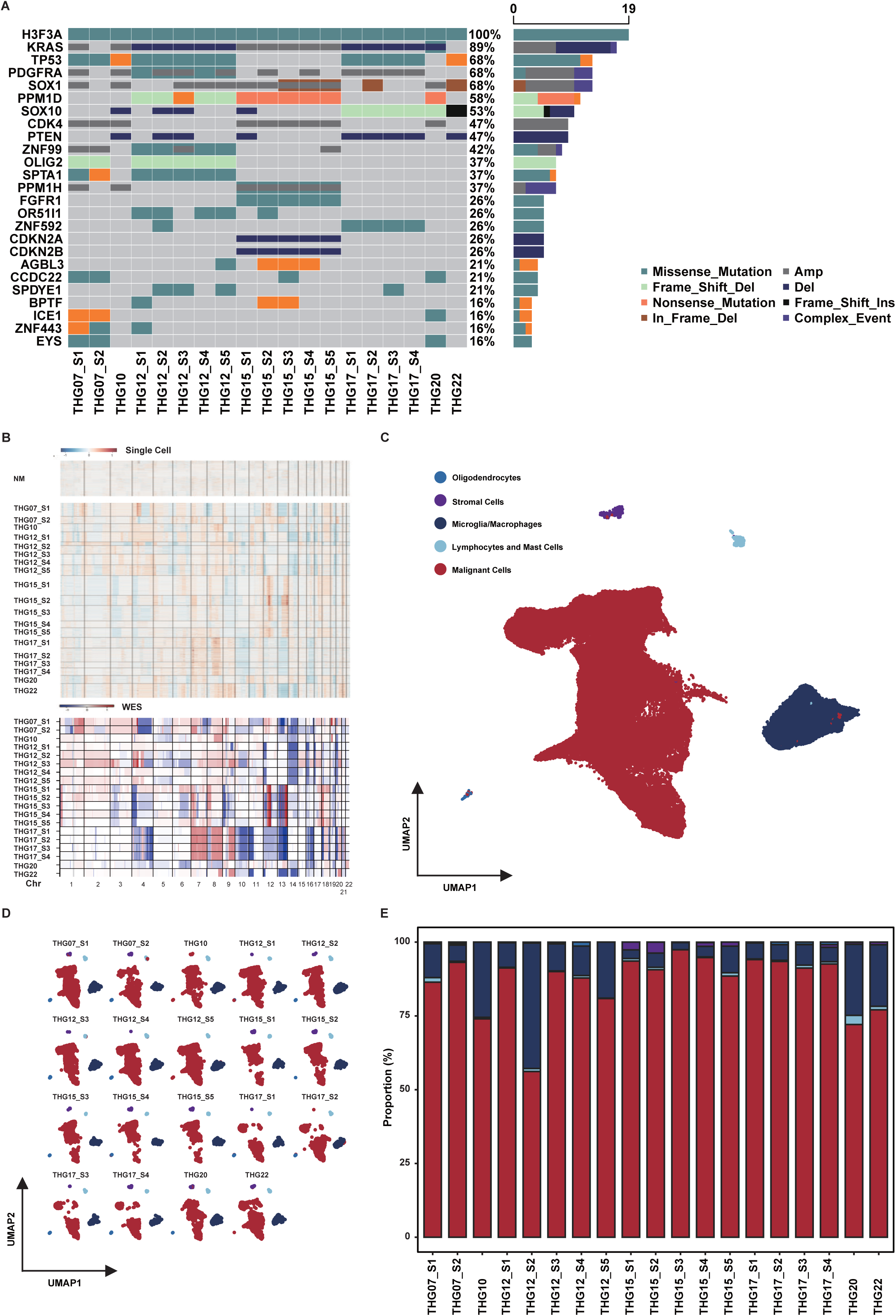
Genomic, transcriptional, and cellular landscapes of spinal cord H3K27-altered DMG. (A) Genomic landscape of 19 samples from 7 patients with spinal cord H3K27-altered DMG. The left panel demonstrates statuses of 20 recurrently altered genes in each tumor sample, color– and pattern-coded by alteration types, with the relative abundance of alterations in each gene shown on the right. Note that when more than one alteration type was present at a given locus in any sample (presented as color overlay on the left), the combined effect was categorized as a “Complex_Event” (purple) in the bar chart on the right, instead of the individual alterations. (B) Copy number variation (CNV) profiles inferred from scRNA-seq (middle panel) and WES (bottom panel) results of spinal cord H3K27-altered DMG are shown. Each row corresponds to malignant cells from each tumor sample, and data within a sample are ordered by the overall CNV patterns. CNV profiles of non-malignant cells (NM) are shown in the top panel for reference. (C) UMAP plot of all single cells pooled from all 19 spinal cord H3K27-altered DMG samples, color-coded by cell types. (D) UMAP plots of single cells from each tumor sample. (E) Percentages of different cell groups in each tumor sample.

We next characterized the cellular composition of all 19 spinal cord H3K27-altered DMG tumor samples using scRNA-seq data (Figure 3B). In total, 119,967 cells passed our stringent quality controls (Supplementary information, Figure S1). We utilized graph-based clustering to group all single cells into one large cluster of tumor cells and four smaller non-malignant clusters. Based on highly expressed markers of specific cell types, the four smaller clusters were further classified into microglia/macrophages (using markers such as CD14 and CD68), lymphocytes and mast cells (e.g., PTPRC), oligodendrocytes (e.g., PLP1), and stromal cells (e.g., RGS5) (Figure 3C; Supplementary information, Figure S2). We then analyzed the copy number variations (CNV) in cells from the large cluster. Cells with CNV patterns that matched those obtained from WES were identified as malignant and subjected to further analysis (Figure 3B, 3C). Accordingly, these analyses resulted in a consistent classification of the sampled cells into malignant and non-malignant subsets. Notably, malignant cells (104,435 in number, 87% of total cells) predominated in all DMG tumor samples (Figure 3D, 3E), which aligns with findings from previous studies [16, 33]. By contrast, microglia and macrophages constituted 11% of all cells (13,349 in number), while immune cells comprised only 0.8% of all cells (969 in number). This is even lower than the previously reported 4% (95 immune cells in 2,458 total cells) [16].

To account for intra-tumoral heterogeneity, we applied non-negative matrix factorization (NMF) to the malignant cells from each tumor sample to identify continuous variability in cellular states [34]. Hierarchical clustering of the NMF patterns based on overlapping gene expression revealed five meta-modules composed of recurrent heterogeneous programs (Figure 4A). The meta-modules consisted of 38–67 genes that frequently appeared across overlapping programs from multiple tumor samples, and each meta-module was present in all 7 patients (Figure 4A). We then denominated the meta-modules according to their component genes and functional enrichment (Figure 4A; Supplementary information, Table S3). Three of the meta-modules were associated with neural differentiation pathways and designated as AC-like (astrocytic differentiation, such as GFAP and APOE), OPC-like (oligodendrocyte precursors, such as PDGFRA and OLIG2) and NPC-like (neural precursors, such as DCX and STMN2), in accordance with previous studies [33]. One meta-module was linked to cell cycling (such as TOP2A and CENPF) and named accordingly. It is worth noting that the meta-module associated with hypoxia and angiogenesis (such as VEGFA and NDUFA4L2) was designated as “hypoxia-related” (Figure 4A). We then classified malignant cells from tumor samples based on the highest-scored “identity” meta-module, which included NPC-like, AC-like, OPC-like, and hypoxia-related modules, while excluding the cell cycle meta-module. We plotted this classification in a two-dimensional cell-state representation map developed by Neftel et al [33] (Figure 4B). The proportions of the four cell states in the malignant cells of each tumor sample are presented in Figure 4C.

**Figure 4.**
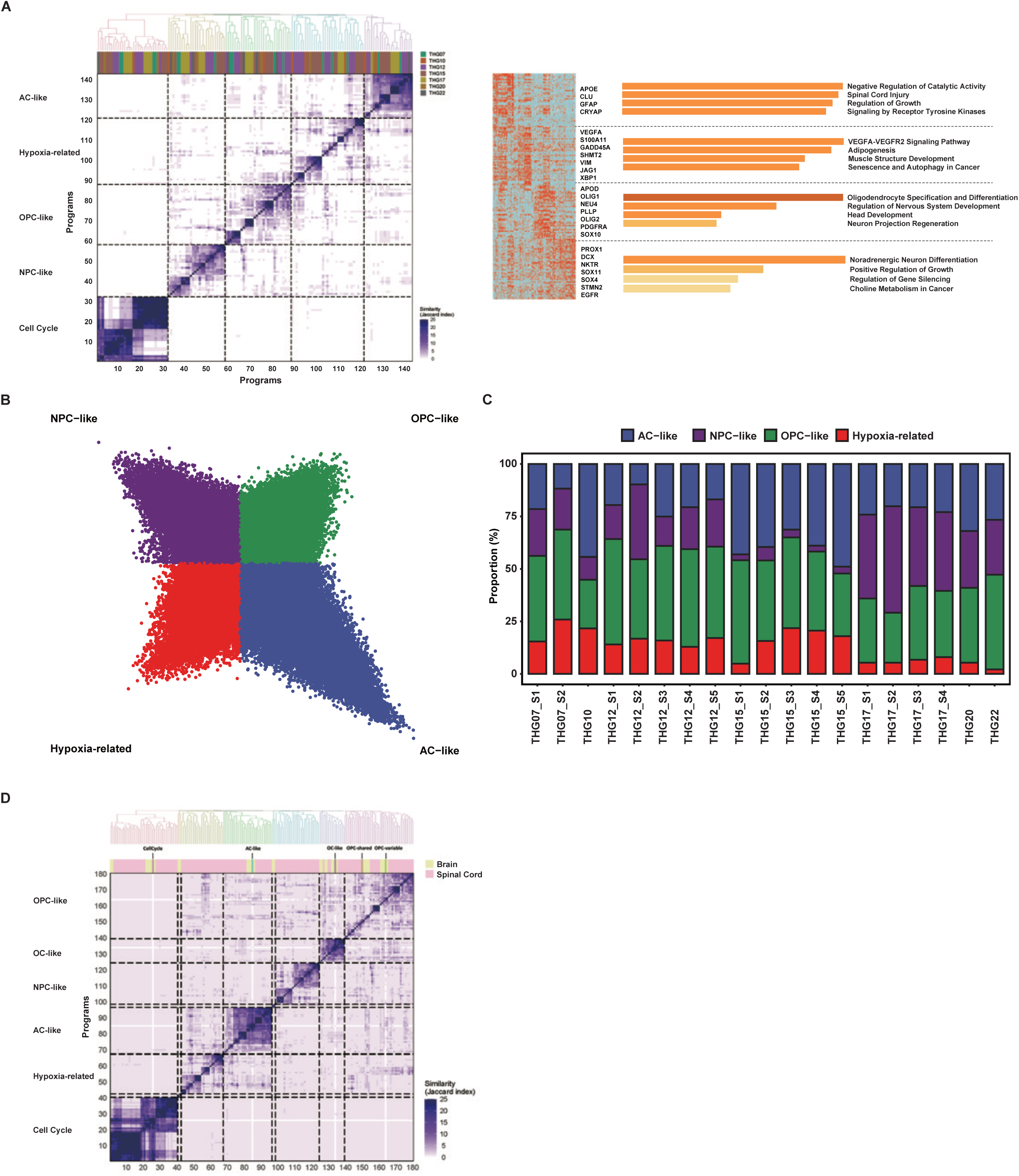
Heterogeneity of malignant cell states in spinal cord H3K27-altered DMG. (A) The main heatmap (bottom) depicts pairwise similarities between 143 non-negative matrix factorization (NMF) programs from all 19 spinal cord H3K27-altered DMG samples, ordered by hierarchical clustering. The top panel shows groups of potential clusters, which were used to define meta-modules. Assignments of programs to patients are presented as color-coded bars on top of the heatmap. Functional enrichment analysis of genes in each meta-module using Metascape is presented on the right, with selected pathways labeled. The results are expressed as −log10 false discovery rate (FDR) adjusted p-values (hypergeometric test). (B) Two-dimensional cell-state representation map embeddings of all 104,435 malignant cells from 19 spinal cord H3K27-altered DMG samples. Each quadrant corresponds to one cellular state, with the exact position of each malignant cell (dot) reflecting its relative scores for the four meta-modules (STAR Methods). (C) Stacked bar plot showing the relative proportions of malignant cells of four cellular states as defined in (B) in each spinal cord H3K27-altered DMG sample. (D) The main heatmap (bottom) depicts pairwise similarities between NMF programs from spinal cord H3K27-altered DMG and published data of brain DMG, ordered by hierarchical clustering. The top panel shows groups of potential clusters. Assignments of programs to tumor locations are presented as color-coded bars on top of the heatmap. Programs defined in the original article of brain DMG (i.e., CellCycle, AC-like, OC-like, OPC-shared, and OPC-variable) are also indicated.

Interestingly, the hypoxia-related meta-module identified in all seven patients with spinal cord H3K27-altered DMG was not reported in the previous study that focused on intracranial DMG [16]. To determine whether this meta-module exists in H3K27-altered DMG in the brain, we reevaluated the expression programs of intracranial DMG using published scRNA-seq data alongside our analysis of intra-tumoral heterogeneity [16]. Hierarchical clustering revealed a high degree of similarity between the three meta-modules defined in the initial research on intracranial H3K27-altered DMG, including OPC-like, AC-like, and OC-like (oligodendrocytic differentiation, such as MBP and PLP1) and those defined in our study. This indicates a consistent gene expression profile of H3K27-altered DMG, regardless of the anatomical location (Figure 4D). However, no expression programs from intracranial DMG could be clustered with the NPC-like or hypoxia-related meta-modules derived from the spinal cord DMG cohort (Figure 4D), suggesting specific expression programs are unique to spinal cord H3K27-altered DMG tumor cells. Interestingly, in patient THG15 with spinal cord DMG who had LM, the expression of hypoxia-related gene expression constituted a significantly smaller proportion in the brain metastasis (THG15_S1) compared to the other four spinal cord tumor samples from the same patient (THG15_S2-S5; Figure 4C), which further supports the specificity of hypoxia-related gene expression program to spinal cord DMG.

To investigate whether there were transitional dynamics among the four cellular states, we conducted RNA velocity analysis. The overall vector field indicated a direction from the hypoxia-related and OPC-like states to the AC-like and NPC-like states (Figure 5A). This suggests a flow from a stem-like phenotype of malignant cells to more differentiated cellular states. The result supports that OPC-like cells have a stem-like nature, consistent with observations in intracranial H3K27-altered DMG [16]. More importantly, this indicates that the hypoxia-related tumor cells specific to spinal cord DMG exhibit inherent stemness.

**Figure 5.**
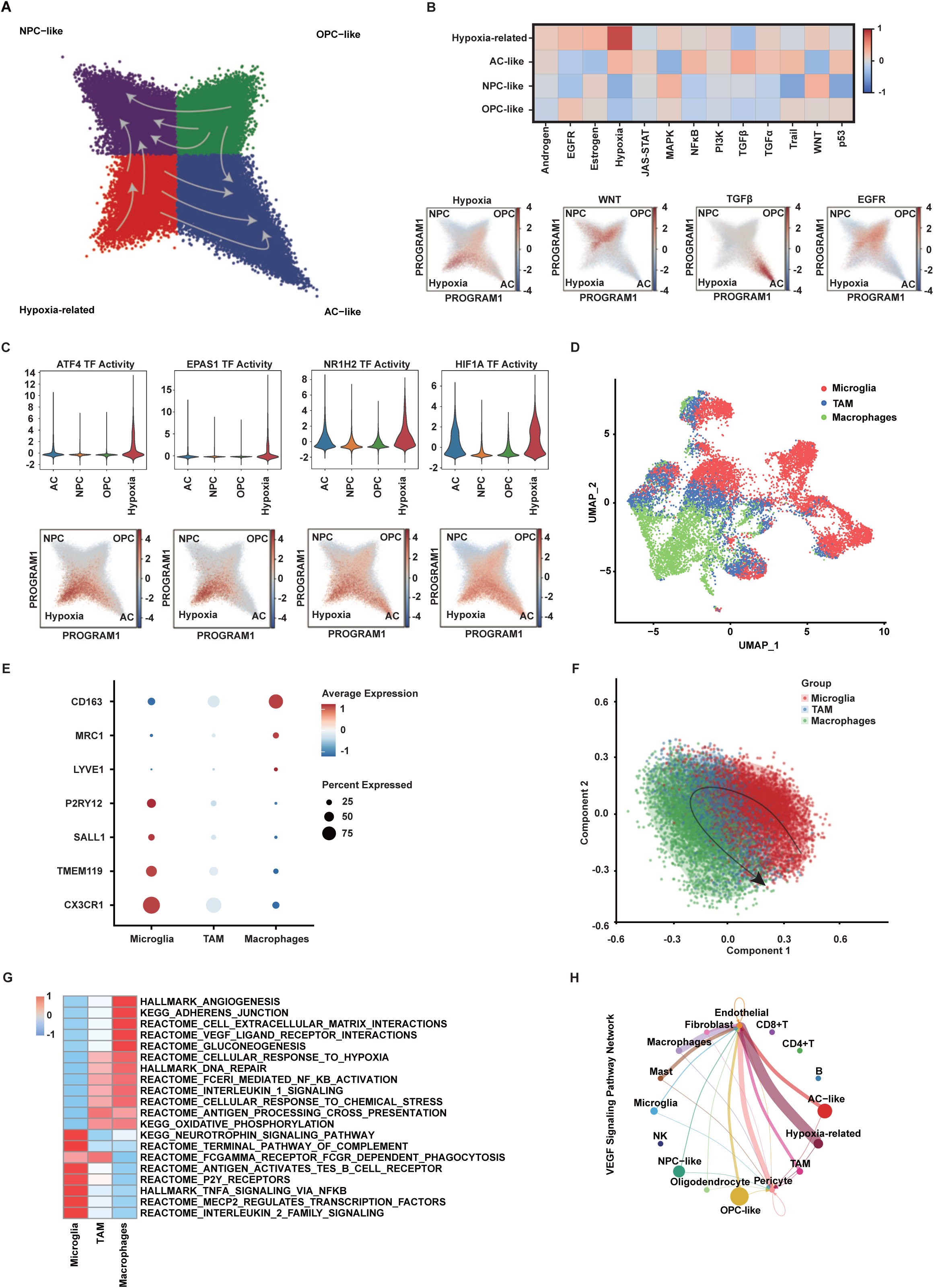
Functional heterogeneity of malignant and tumor associated myeloid cells in spinal cord H3K27-altered DMG. (A) The directional flow of the velocity field per RNA velocity analysis is superimposed on the two-dimensional cell-state representation map embeddings of all malignant cells. (B) Relative activities of 13 selected pathways in malignant cells of each state are shown as the collective mean in the top panel. The bottom panel shows four two-dimensional cell-state representation maps embeddings of all malignant cells, as in Figure 4B, with the color of each cell reflecting its relative activity of the indicated pathway. The selected pathways represent the most highly activated pathway for each cell state (i.e., hypoxia for hypoxia-related cells, WNT for NPC-like cells, TGFβ for AC-like cells, and EGFR for OPC-like cells). (C) In the top panel are violin plots showing activity levels of genes encoding transcription factors involved in hypoxia pathway including ATF4, EPAS1, NR1H2 and HIF1A in malignant cells of each state. Activities of these transcription factors in malignant cells are also reflected on two-dimensional cell-state representation maps in the bottom panel. (D) UMAP plot of myeloid cells pooled from all 19 spinal cord H3K27-altered DMG samples, color-coded by cell types. Tumor associated microglia/macrophages (TAM) represent an intermediate myeloid cell state with concurrent expression of signature genes of both microglia and macrophages. (E) Expression levels of signature genes of microglia (e.g., CX3CR1 and P2RY12) and macrophages (e.g., CD163 and MRC1). Percentage of cells in each group expressing indicated genes is also reflected by the sizes of dots. (F) Inferred developmental trajectory among tumor associated myeloid cells by SCORPIUS, with the arrow indicating direction of trajectory. The cells are presented on a PCA plot based on scores of the first two principal components (See Supplementary information, Table S5), and are color-coded by myeloid cell subsets. (G) Relative activities of selected pathways in each subset of myeloid cells are shown as scale scores based on GSVA analysis. The selected pathways represent significantly upregulated pathways in each myeloid cell subset. (H) Chord diagram showing predicted cell-to-cell interactions of VEGF signaling pathway network in spinal cord H3K27-altered DMG.

To explore the functional statuses of malignant cells in various cellular states, we performed PROGENy analysis to infer pathway activities. We found that different cellular states corresponded with the activation of distinct pathways (Figure 5B; Supplementary information, Table S4). Notably, the hypoxia signaling pathway was highly activated in cells related to hypoxia as expected, while AC-like cells displayed activation in NF-κB and TGFβ signaling pathways. In contrast, NPC– and OPC-like cells demonstrated activated WNT and EGFR signaling pathways, respectively. Additionally, we used DoRothEA for transcription factor inference, which revealed highly activated transcription factors implicated in hypoxia signaling pathways such as ATF4, EPAS1, NR1H2, and HIF1A in hypoxia-related cells (Figure 5C; Supplementary information, Figure S3A, Table S4), corroborating pathway activation. Similarly, distinct transcription factor expression profiles were also observed in AC-, NPC-, and OPC-like cells (Supplementary information, Figure S3A), aligning with our pathway inference results.

### State transition of tumor-associated microglia to macrophages reflects enrichment of hypoxia-related gene programs and functional pathways

The tumor microenvironment (TME) is crucial in shaping tumor cell behavior. We hypothesized that the activation of the hypoxia-related signaling pathway reflects how spinal cord DMG tumor cells respond to a relatively hypoxic TME and that a similar response may also be seen in non-malignant cells within the TME. To characterize the TME, we profiled the non-malignant cell clusters, including immune and stromal cells shown in Figure 3C, using subclustering analysis. We identified CD8+ T cells, CD4+ T cells, mast cells, B cells, and NK cells in the immune cluster (Supplementary information, Figure S3B, S3C). In the stromal cluster, we identified pericytes, endothelial cells, and fibroblast cells (Supplementary information, Figure S3D, S3E). Overall, our results revealed few tumor-infiltrating lymphocytes (of less than 1% of total cells; See also Figure 3C), supporting the notion of the immunologically “cold” nature of spinal cord DMG [35].

Next, we used principal component analysis (PCA) to evaluate the diversity within the microglia/macrophage cluster, which was the most prominent subset of non-malignant cells in the TME (Supplementary information, Figure S3F). In accordance with previous reports [36], the second principal component reflected an inflammatory program (Supplementary information, Table S5). Conversely, the principal component 1 (PC1) score reflected two mutually opposing programs: One with higher expression was associated with macrophages (PC1-high, characterized by markers such as CD163 and TGFBI) and the other with lower expression was associated with microglia (PC1-low, represented by markers such as CX3CR1 and P2RY12) (Supplementary information, Table S5). Consequently, we classified PC1-high cells as infiltrating macrophages and PC1-low cells as spinal cord-resident microglia (Supplementary information, Figure S3F). Interestingly, a subset of cells from the microglia/macrophage cluster expressed both microglia markers (such as CX3CR1 and P2RY12) and macrophage markers (such as CD163 and MRC1) (Figure 5D, 5E; Supplementary information, Figure S3G). This finding supports a continuum in gene expression between microglia and macrophages in H3K27-altered DMG, consistent with reports regarding IDH-mutant gliomas [35]. We defined the intermediate state of microglia/macrophages that expressed both markers as tumor-associated microglia/macrophages (TAM) (Supplementary information, Figure S3F, S3G). We then conducted trajectory inference analysis, which revealed a developmental trajectory from microglia through TAM to macrophages (Figure 5F). This suggests a state transition and remodeling of microglia within the TME.

Analysis of the differentially expressed genes among the three subsets of microglia/macrophages revealed that microglia had high expression of TNF and IL1B, while macrophages exhibited increased expression of VEGFA and MMP14 (Supplementary information, Figure S3H). In comparison, TAM exhibited higher expression of the complement gene C1QC and several antigen-presentation-related genes in the HLA family (Supplementary information, Figure S3H, S3I, S4). Further evaluation of pathway activity in the three subsets using gene set variation analysis (GSVA) revealed that microglia were strongly enriched in the TNFα/NF-κB, IL2, and neurotrophin signaling pathways (Figure 5G). In comparison, macrophages exhibited enrichment in pathways related to VEGF ligand interactions, angiogenesis, and cellular response to hypoxia. TAM not only showed an enrichment in antigen processing and presentation pathway activity but also exhibited repressed activities in pathways enriched in microglia, such as neurotrophin signaling, while displaying increased activity in macrophage-enriched pathways like the cellular response to hypoxia (Figure 5G). Interestingly, in patient THG15, the proportion of macrophages among myeloid cells was significantly lower in brain metastasis (THG15_S1) compared with spinal lesions (THG15_S2-S5; Figure S5), which paralleled the observation for hypoxia-related malignant cells (Figure 4C). Together, these results support our earlier inference that TAM represents a transitional state from microglia to macrophages, suggesting that this transition may reflect cellular responses to hypoxia with increasing VEGF activity and angiogenesis.

### Intra-tumoral functional heterogeneity and extensive VEGF interaction network in TME

Results from the above functional analyses implied that hypoxia may play a role in sculpting both tumor cells and the TME in H3K27-altered DMG of the spinal cord. We thus hypothesized that extensive angiogenic-related cell-to-cell interactions occur within the TME. We performed a comprehensive cell-to-cell interaction analysis that included all identified cell types so far (Supplementary information, Figure S6A). We found that the VEGF signaling pathway network was present between most cell types except CD8+T cells, CD4+T cells, B cells, and NK cells (Figure 5H). Endothelial cells and pericytes acted as the core in the predicted networks, exhibiting the highest number of interactions with other cell types. Hypoxia-related tumor cells had the strongest interaction with endothelial cells. Additionally, microglia, TAM, and macrophages exhibited a progressive increase in interaction strength with endothelial cells through the VEGF pathway network (Figure 5H), which was further supported by GSVA results (Figure 5G). Other cell-to-cell interaction networks also corroborated results from functional analyses (Supplementary information, Figure S6B). For instance, microglia were the pivot of the TNF signaling pathway, implying their anti-tumor function as the primary immune cells in the central nervous system. Different subsets of malignant cells played distinct roles in interaction networks (Supplementary information, Figure S6B). Specifically, AC-like cells were the only malignant cells involved in TGFβ and periostin signaling pathway networks. The latter, which is inducible by hypoxia, is associated with recruiting TAM and promoting glioma progression [37–39]. NPC-like cells represented the center of the ncWNT and EGF signaling pathway networks, whereas OPC-like cells played a crucial role in the MPZ signaling pathway network. Collectively, our data suggest that hypoxia-related malignant cells and macrophages constitute the key cellular components in the VEGF interaction network in spinal cord H3K27-altered DMG. Moreover, different states of malignant cells may display distinct functions through highly diverse gene program expression and varied network interactions with other cellular components of the TME.

To investigate whether the cellular architecture varied across different spatial contexts within the same tumor, we compared the proportions of malignant and myeloid cells in samples THG12, THG15, and THG17, where multiple sampling was performed. THG12_S2, THG15_S3, and THG17_S1 represented tumor core tissues, while the remainder were obtained from the periphery (Table 1, Supplementary information Figure S5A). Interestingly, we did not observe any consistent trends for specific cell types across core versus peripheral tissues, suggesting compositional variability in spinal cord H3K27-altered DMG (Supplementary information, Figure S5B, C). However, it is worth noting that a significantly increased proportion of total myeloid cells was found in the core sample in THG12 (THG12_S2) compared with the peripheral samples or any other sample tested (Supplementary information, Figure S5B), suggesting marked inter– and intra-tumoral heterogeneity in the cellular signature of spinal cord DMG.

### Patient-derived spinal cord H3K27-altered DMG cells are sensitive to combined HDAC and EZH2 inhibition

Given the key role of the oncohistone H3K27M in the tumorigenesis of DMG through epigenetic reprogramming, therapies targeting epigenetic reprogramming are considered promising treatment strategies with accumulating preclinical and clinical evidence of efficacy [40–45]. We have observed promising responses in recurrent H3K27-altered DMG patients treated with a combination of panobinostat, a histone deacetylase (HDAC) inhibitor, tazemetostat, an EZH2 inhibitor, and immunotherapy [7]. To investigate changes in expression profiles of the four malignant cell state programs during treatment, we isolated spinal cord DMG tumor cells from patient-derived surgical tissues (THG15_S3; Figure 6A). Protein expression of signature genes of each cell state (VEGFA, GFAP, PDGFRA, and SOX11 for hypoxia-related, AC-like, OPC-like, and NPC-like, respectively) at baseline was validated with immunofluorescence (Figure 6B). Consistent with our previous report, primary cultures of patient-derived spinal cord H3K27-altered DMG cells demonstrated sensitivity to the combined treatment of panobinostat and tazemetostat (Figure 6C). This combination was shown to further reduce global H3K27 trimethylation while increasing H3K27 acetylation [7]. Concurrently, RNA sequencing analysis revealed sustained expression repression of the majority of feature genes in the hypoxia-related meta-module, including VEGFA, S100A11, GADD45A, and SHMT2 (Figure 6D; See also Supplementary information, Table S3), all of which are known to promote cellular proliferation in high-grade gliomas [46–49]. A corresponding reduction in protein expression was also observed through western blotting analysis (Figure 6E). Collectively, these results suggest that combined inhibition of HDAC and EZH2 may effectively suppress the proliferation of spinal cord H3K27-altered DMG cells through downregulating the expression of genes associated with the hypoxia-related gene program.

**Figure 6.**
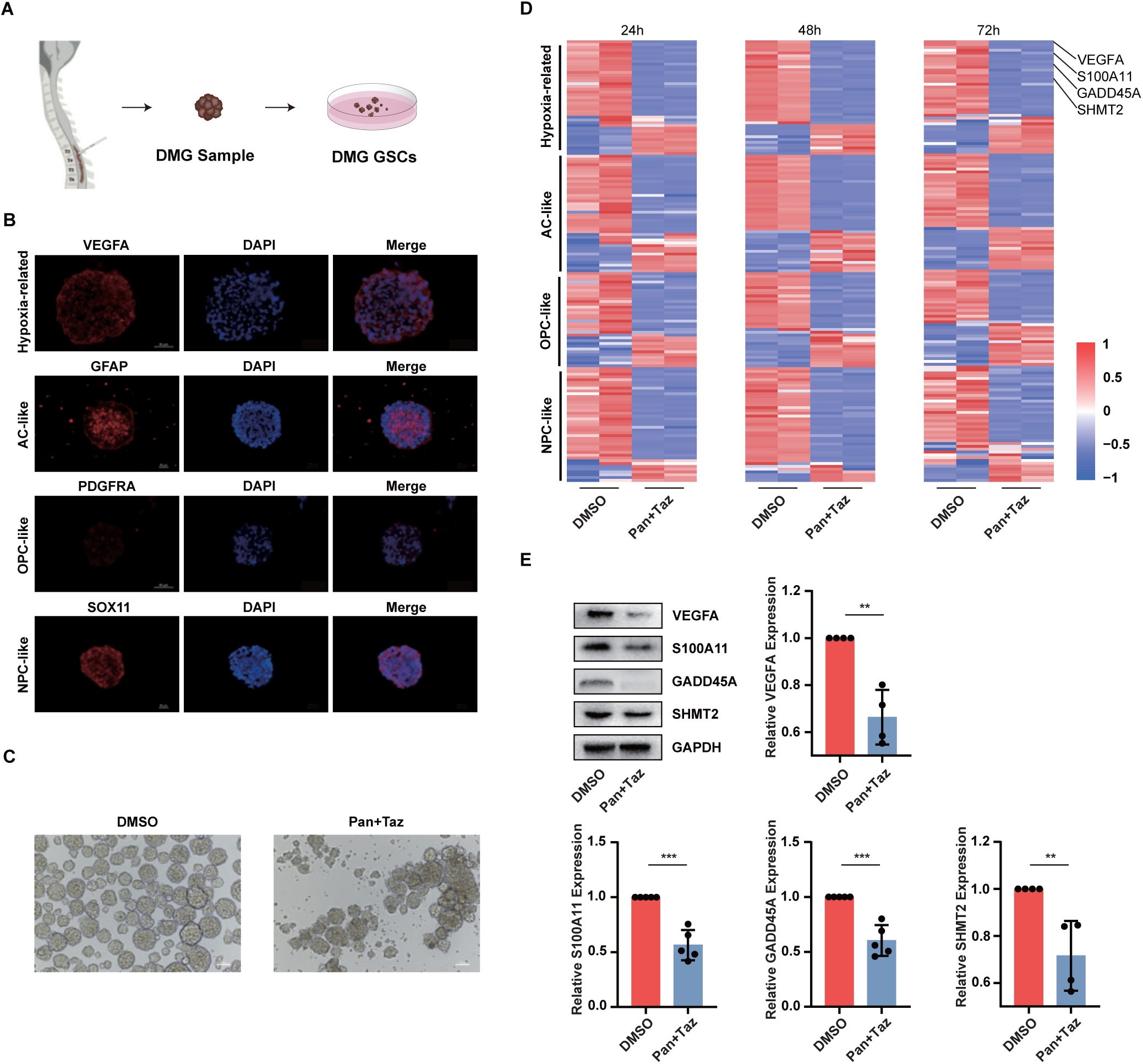
Combined epigenetic agents suppressed hypoxia-related gene expression and inhibited growth of spinal cord H3K27-altered DMG cells in vitro. (A) Schematic of workflow for generating patient-derived spinal cord H3K27-altered DMG cell cultures. (B) Micrographs of immunofluorescence in spinal cord H3K27-altered DMG neurospheres stained for the indicated marker of each meta-module (red) and DAPI (blue). (C) Response of DMG neurospheres to combined panobinostat (Pan) and tazemetostat (Taz) or DMSO control after incubation for 72 hours. (D) Relative expression of feature genes of each meta-module in DMG neurospheres after 24, 48, and 72 hours of incubation with DMSO control or combined panobinostat (Pan) and tazemetostat (Taz). Results are shown in columns of duplicates. (E) Western blots of VEGFA, S100A11, GADD45A, SHMT2, and GAPDH in DMG neurospheres after incubation with combined epigenetic agents or DMSO for 48 hours are shown in the top left panel. The respective protein levels in epigenetic-agent-treated cells relative to DMSO control are also shown. **, p<0.01, ***, p<0.001, Student’s t test.

## Discussion

We investigated intra– and inter-tumoral heterogeneity in spinal cord H3K27-altered DMG through single-cell transcriptomic and genomic analyses and established putative cellular hierarchies. We identified a hypoxia-related gene meta-module that was consistently expressed in spinal cord DMG, but entirely absent in intracranial H3K27-altered DMG. Malignant cells with differential expression of the hypoxia-related gene program played a central role in a TME characterized by extensive activation of angiogenic and VEGF pathway networks. Using RNA velocity analysis, we inferred that hypoxia-related tumor cells possess a stem-like phenotype, similar to OPC-like tumor cells. Significantly, epigenetic treatment of patient-derived DMG cultures with combined inhibition of HDAC and EZH2 suppressed hypoxia-related gene expression and inhibited tumor cell proliferation in vitro. Combined with our previous clinical study that showed efficacy of applying HDAC and EZH2 inhibitor-based immunotherapy in treating DMG patients [7], these findings suggest the underlying mechanism for this promising therapeutic regimen in spinal cord H3K27-altered DMG.

Prior to this work, few studies have examined the genetic and transcriptional signatures of spinal cord H3K27-altered DMG [17], partly attributable to its rare occurrence. Our study amassed WES and scRNA seq data from 7 patients and 19 tumor samples, spanning primary, recurrent, and metastatic lesions (Table 1), representing the largest cohort in a single study to date with a transcriptional landscape at single-cell resolution. We observed a high prevalence of TP53 mutation, in addition to the DMG-defining H3F3A mutation, and mutation and/or amplification of PDGFRA (Figure 3A). These genetic alterations are consistent with findings in previous reports on H3K27-altered DMG of the spinal cord [50–52] or brain [10, 17, 32, 53]. Moreover, we also discovered frequent deletion in PTEN (Figure 3A), which was not observed in spinal cord DMG in prior studies [17, 32]. Together, these results support consistent albeit diverse patterns in the mutational programs driving tumorigenesis in DMG regardless of tumor location [8, 54]. Additionally, in line with previous studies of primarily intracranial tumors [16, 17], malignant cells dominated the spinal cord H3K27-altered DMG landscape, while non-malignant cells primarily consisted of microglia and macrophages, with a striking paucity of tumor-infiltrating lymphocytes (Figure 3C; Supplementary information, Figure S6A), suggesting an immunologically “cold” tumor with clinically consequential implications [9]. Such immune-cold tumors have a limited response to immunotherapy, as demonstrated by the failure of PD-1/PD-L1 blockade to improve overall survival in high-grade glioma patients, including DMG [55–58], underscoring the need for combinatorial strategies in clinical settings. Collectively, these results reinforce common oncogenic drivers and the immune-cold microenvironment as key characteristics of H3K27-altered DMG, transcending anatomical locations.

Notably, compared to the brain, the spinal cord has unique anatomical constraints, featuring a narrow, tubular, rigid vertebral canal with limited compensatory space and exhibiting distinct vascular characteristics, including significantly lower vascular density and differential perfusion patterns [59, 60]. Based on differential expression of recurrent gene programs, we identified four molecularly defined cellular states in spinal cord H3K27-altered DMG: OPC-like, AC-like, NPC-like, and hypoxia-related, with each characterized by distinct functional pathway activation and cell-to-cell interactions (Figure 4A; Supplementary information, Figure S6B). Similar to intracranial H3K27-altered DMG [16], OPC-like malignant cells in spinal cord DMG exhibited inferred stemness and plasticity (Figure 5A). Surprisingly, developmental trajectory inference indicated that hypoxia-related malignant cells may also contribute as a source of stem cells in spinal cord H3K27-altered DMG (Figure 5A). Compared to the mesenchymal-like (MES-like) signature defined in a previous study of primarily intracranial DMG that also expressed genes such as GAP43 and DDIT3 [17], the hypoxia-related signature in spinal cord DMG differed through accentuated expression of angiogenic factors, including VEGFA and JAG1 (Figure 4A). Importantly, we confirmed the absence of the hypoxia-related gene program in published data on pediatric pontine and thalamic DMG (Figure 4D). In line with this, in patient THG15, intracranial metastases from spinal cord DMG comprised significantly fewer hypoxia-related malignant cells than the tumor within the spinal cord (Figure 4C). Together, these findings indicate the presence of a unique group of hypoxia-related tumor cells in DMG of the spinal cord with a potential role as glioma stem cells and support variability in cellular architecture and transcriptomic landscape in DMG across spatial contexts despite the common H3K27 alteration.

Hypoxia has long been associated with promoting malignant behaviors of tumor cells and cancer progression, leading to a worsened prognosis [61, 62]. For instance, a hypoxic tumor microenvironment could alter lactate metabolism and stimulate angiogenesis through upregulating VEGF in spinal cord malignancies[63]. Interestingly, despite the unique presence of hypoxia-related tumor cells, spinal cord H3K27-altered DMG patients even survived significantly longer than thalamic and pontine DMG patients (Figure 2). The underlying reasons could be multifactorial. Firstly, the patient’s survival may be confounded by the location involved, extrinsic to tumor behaviors. The thalamus and brainstem are crucial to critical life functions, and their involvement by either benign (e.g., hemangioblastoma) or malignant (e.g., high-grade gliomas) pathologies has been associated with a poorer prognosis compared with participation of the spinal cord [64, 65]. Secondly, the hypoxia-related tumor cells in spinal cord DMG may adopt a less malignant phenotype that contributes to improved survival in patients. The hypoxia signature has been implicated in MES-like malignant cells in glioblastoma [33, 66], which is in contrast to an astrocyte-related signature in MES-like cells in intracranial H3K27-altered DMG [16, 67]. Similarly, NPC-like cells, absent in intracranial H3K27-altered DMG (Figure 4B and D), have been found in abundance and actively proliferating in glioblastoma with an expression signature consistent with what was defined here (e.g., STMN2) [33]. These observations may point to an underrecognized similarity in cellular states between DMG of the spinal cord and glioblastoma. As MES-like cells in glioblastoma have the least proliferative activity when compared with OPC-like, NPC-like, and AC-like cells [33, 66, 68], the hypoxia-related cells in spinal cord H3K27-altered DMG, with a partly similar transcriptomic signature, may also be in a relatively quiescent state despite stemness potential [69]. Taking into account that MES-like glioblastoma cells lack consistent VEGFA expression [67], which is highly enriched in hypoxia-related cells in spinal cord DMG, further in vitro and in vivo studies on the functions of hypoxia-related DMG cells are needed to better gauge their biological and clinical implications and help dissect heterogeneity in tumorigenesis and biological behaviors of high-grade gliomas.

Myeloid cells, including microglia and macrophages, represent the most significant immune cell subpopulation in high-grade gliomas, including DMG [9, 35], as corroborated here (Figure 3C, Supplementary information, Figure S5B). We found a state transition from microglia-like to macrophage-like phenotype in tumor-associated myeloid cells with increasing activities of angiogenic signaling pathways and involvement in the VEGF signaling network (Figure 5G, 5H), which suggests the promotion of angiogenesis by DMG-infiltrating macrophages. A similar association between macrophage-like state polarization and histologically high-grade hallmarks such as angiogenesis has been implicated in IDH-mutant gliomas [36]. Since expression profiles of myeloid cells may be altered by the global glioma TME [36], concurrently activated hypoxia-related signaling pathways in hypoxia-related malignant cells and macrophages, without a direct VEGF signaling network interaction (Figure 5H), may reflect a globally hypoxic TME in spinal cord H3K27-altered DMG. This is also supported by an association between the hypoxia-related MES-like signature in malignant cells and increased macrophage-to-microglia ratio in other high-grade gliomas [67]. Recently observed age-dependent variation whereby macrophages predominated in adult DMG and microglia in pediatric DMG, linked to differences in the proportion of MES-like signature in malignant cells [17], suggests the presence of a tumor-extrinsic impact on myeloid cell polarization in H3K27-altered DMG that may also contribute to sculpting DMG immune TME.

Significant inter– and intra-tumoral heterogeneity in cellular composition was observed in spinal cord H3K27-altered DMG (Supplementary information, Figure S5B, C), which has also been reported in other high-grade gliomas [17, 70]. One notable example was the profuse myeloid cell infiltration in THG12_S2 compared to all other specimens (Supplementary information, Figure S5B). This cannot be mainly explained by the unique treatment history in the patient (e.g., with ONC201; See Table 1 and ref.7 supplemental Fig. S14) given the lower myeloid infiltration in the peripheral samples from the same tumor (THG12_S3 and S5), due to the lack of pretreatment samples. Instead, the marked cellular variability in spinal cord DMG is more likely driven by the diverse mutational programs and heterogeneous CNV even within the same tumor (Figure 3A, B), which may also be responsible for treatment resistance and tumor recurrence [70, 71]. Consequently, multifocal tumor sampling could capture the dynamic heterogeneity of DMG. By comparative analysis of pre– and post-treatment multifocal biopsies, we could further delineate therapy-induced clonal evolution, facilitating the development of dynamic precision therapeutic strategies tailored to tumor adaptation.

Surgically, decompressive laminectomy is recommended for spinal cord DMG primarily for two reasons: Firstly, it expands the spinal canal volume, reduces mass effect and improves spinal cord perfusion, thereby alleviating tumor-induced hypoxia, remodeling TME and its related tumor progression and/or metastasis. Secondly, it decreases postoperative steroid (e.g., dexamethasone, methylprednisolone) dependency-induced immunosuppression, thus synergizing with combination immunotherapy regimens. Beyond surgery, precise therapeutic options are still lacking in H3K27-altered DMG [7, 71]. Various tumorigenic pathways converge on the VEGF axis in DMG [72, 73]. However, VEGF-A blockade with bevacizumab may produce prompt clinical and radiographic responses, primarily by reducing tumor-associated edema through vascular normalization in DMG patients, but fails to demonstrate a conclusive overall survival benefit [74–78], underscoring the need for further combinatorial therapies. We have reported promising therapeutic effects of epigenetic agent-based immunotherapy in spinal cord H3K27-altered DMG patients [7]. Mechanistically, global modulation of histone methylation/acetylation balance may interfere with or revert tumorigenic cascade effects of H3K27M mutation [3, 40, 45, 79], although HDAC inhibitors as a standalone might not be adequate to improve survival in high-grade glioma patients [80, 81] and clinical experience with EZH2 inhibitors in DMG patients is scarce [7]. Here, we showed that combined HDAC and EZH2 inhibition effectively suppressed hypoxia-related gene expression, including VEGFA, and inhibited DMG cell proliferation in vitro (Figure 6C–E), supporting the therapeutic implication thereof. Further studies are required to investigate whether VEGFA blockade could enhance the efficacy of HDAC and EZH2 inhibition-based immunotherapy and the underlying synergistic mechanisms in spinal cord DMG.

Collectively, our study reveals a crucial anatomical dimension that DMG exhibits a location-specific genomic and transcriptional landscape despite shared H3K27 alterations. These findings suggest that primary tumor niches dictate treatment vulnerability, underscoring the need for anatomically guided therapeutic strategies. Additional samples from thalamic and brainstem DMG are warranted for further validation. Significantly, this paradigm — “Targeting molecular mutations/alterations based on anatomical coordinates” — may extend beyond DMG to other molecularly defined yet anatomically divergent malignancies, including: IDH mutant gliomas, BRAF V600E cancers, NTRK-fusion tumors, INI1-deficient tumors, and PAX3-FOXO1+ alveolar rhabdomyosarcomas. Therefore, it could revolutionize precision oncology by delivering therapies tailored to both molecular drivers and their anatomical microenvironment.

## Acknowledgments

We thank Dr. James Jin Wang for his academic support, encouragement, and inspiration. We thank Prof. Da Mi, Yi Lin and Kun Li from School of Life Sciences, Prof. Tienyin Wong from School of Medicine, Tsinghua University, and Prof. Fan Bai from School of Life Sciences, Peking University, for their constructive suggestions. We thank Dr. Yafang Deng from School of Medicine, Tsinghua University, and Benqi Zhao from Department of Radiology, Beijing Tsinghua Changgung Hospital, for their assistance in data analysis.

## Funding

This work was supported by Institute of Biomedicine and Tsinghua-Peking Joint Center for Life Sciences.

## Conflict of interest statement

The authors declare no conflict of interest.

## Authorship statement

All authors were involved in the conceptualization and initiation of the study. Yiwei Xiao, Qiang Gao, Zhihong Qian, Mengyao Li, Yukui Shang, Linkai Jing, Zili Zhen, and Yong Ai were responsible for the study investigation and acquisition of the data. Kehkooi Kee, Guihuai Wang, and Wei Zhang provided pertinent resources and guidance on conduct of the study. Yiwei Xiao, Qiang Gao, and Zhihong Qian drafted the original manuscript, and all authors were responsible for revision of the manuscript. Wei Zhang was responsible for funding acquisition.

## Data and software availability

The datasets generated during the current study are available from the corresponding authors on reasonable request. The high-throughput data have been deposited to the (It will be obtained when we upload the files)

## Figure legends

**Figure S1.**
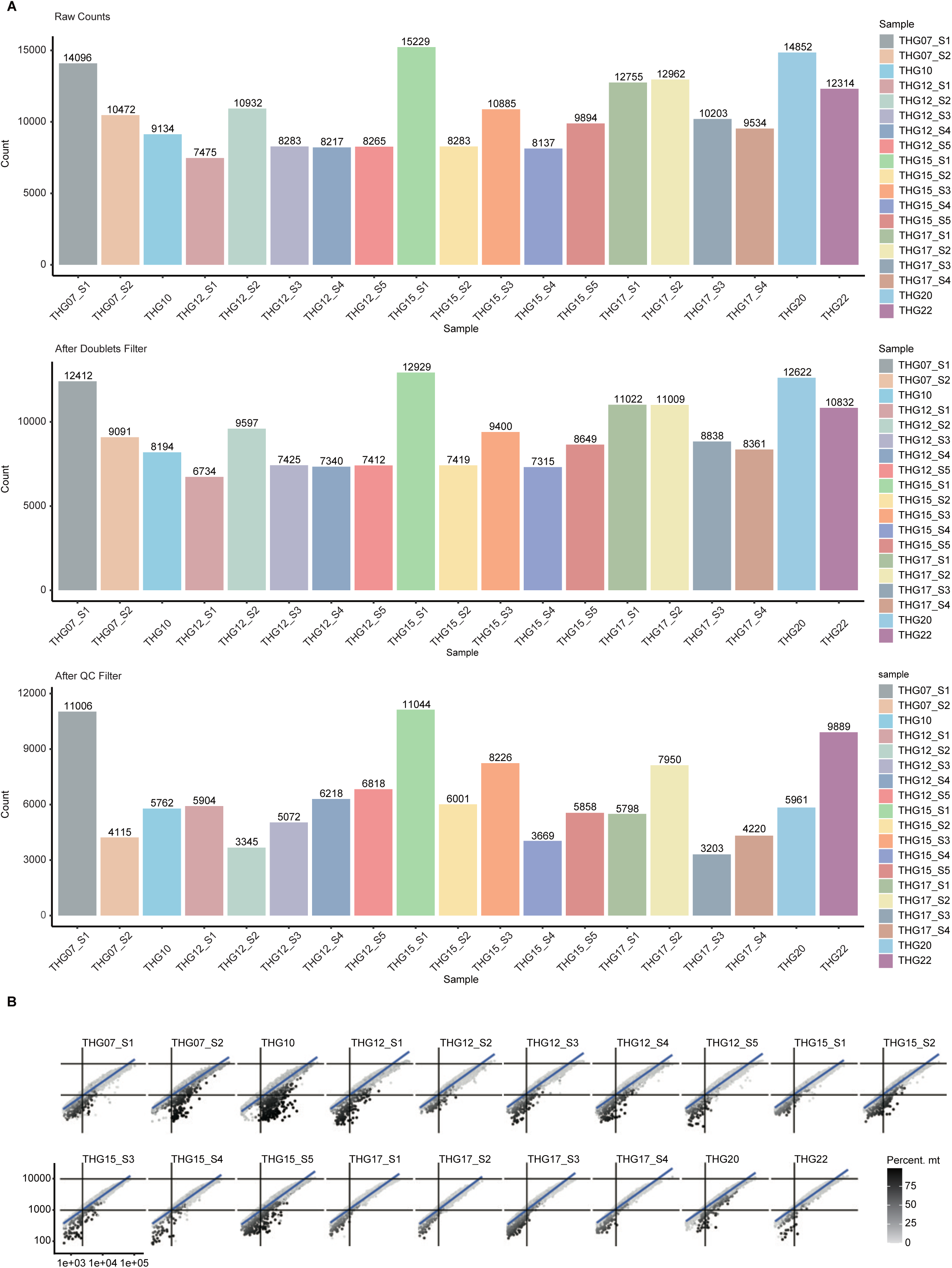
(A) Cells filtered in quality control of scRNA-seq dataset. Top panel: cell counts in raw data; middle panel: cell counts in each sample after doublets filter; bottom panel: cell counts in each sample after all quality control filters. (B) Summary of the quality of each cell in the raw scRNA-seq dataset in each sample. The x-axis represents the captured total RNA counts, the y-axis represents the captured number of genes, and the color intensity represents the proportion of mitochondrial genes in the cellular RNA read counts.

**Figure S2.**
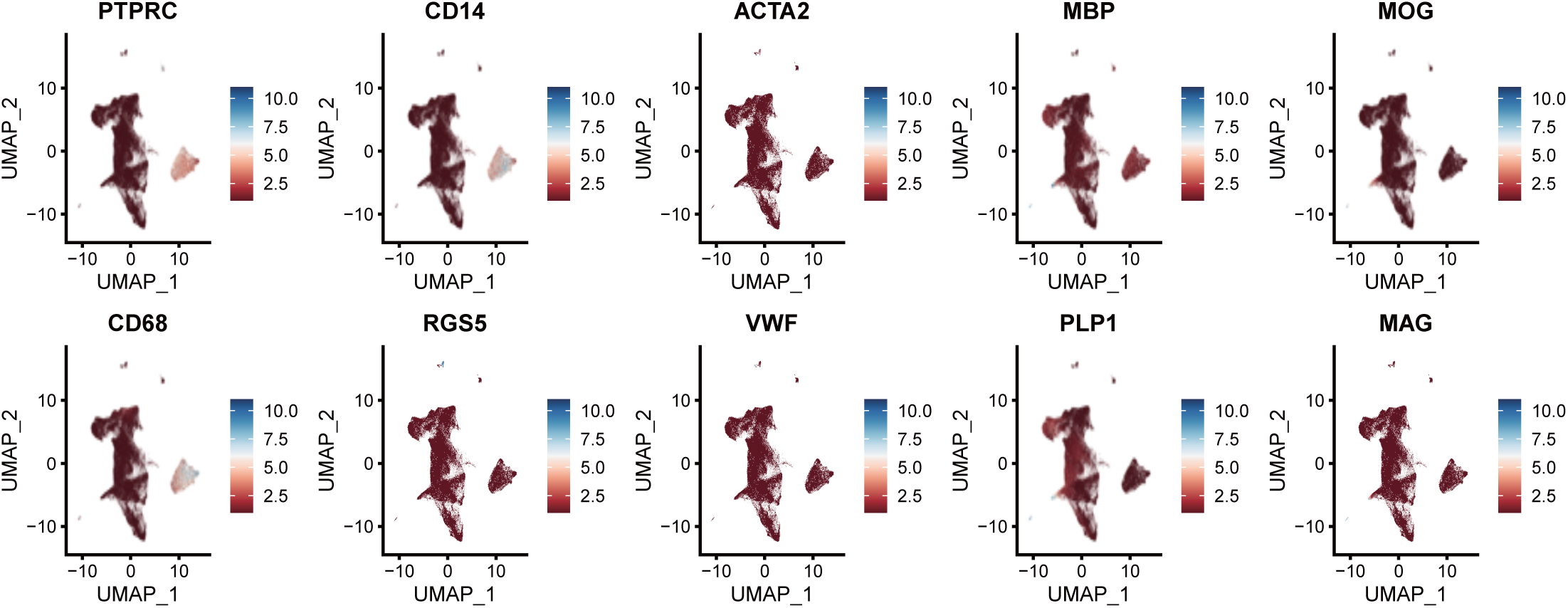
UMAP plots showing expression levels of selected genes in major cell clusters from spinal cord H3K27-altered DMG samples.

**Figure S3.**
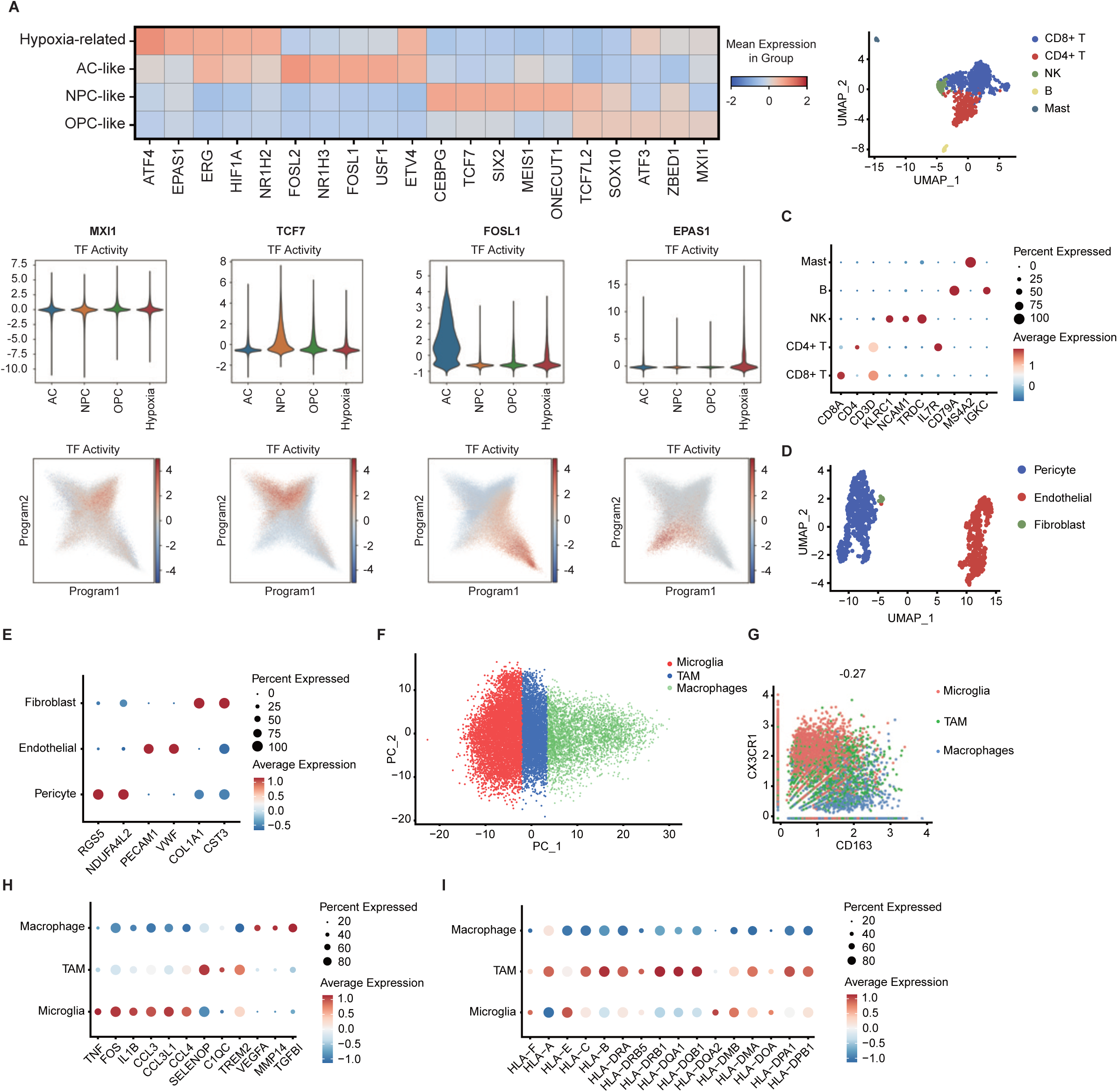
(A) The upper panel shows the expression levels of the top five most activated transcription factors in each of the four malignant cell states. The middle panel shows the relative activities of selected transcription factors in each malignant cell state as violin plots. The bottom panel shows two-dimensional cell-state representation maps embeddings of all malignant cells, with the color of each cell reflecting the relative activity of the indicated transcription factor. (B) UMAP plot of cells from the immune cell cluster. Cells are colored based on the immune cell subsets. (C) Dot plot showing classic marker genes across immune cell subsets. Dot size is proportional to the fraction of cells expressing specific genes. Color intensity corresponds to the relative expression of specific genes. (D) UMAP plot of cells from the stromal cluster. Cells are colored based on the stromal subsets. (E) Dot plot showing classic marker genes across stromal subsets. Dot size is proportional to the fraction of cells expressing specific genes. Color intensity corresponds to the relative expression of specific genes. (F) PCA plot showing the myeloid cells in spinal cord H3K27-altered DMG. (G) Scatter plot showing CX3CR1 and CD163 levels in myeloid cells. Cells are colored by microglia/macrophages subsets. Pearson correlation between the two features is displayed above the plot. (H) Dot plot showing expression of selected differential genes across the microglia/macrophages subsets. The dot size is proportional to the fraction of cells expressing specific genes. The color intensity corresponds to the relative expression of specific genes. (I) Dot plot showing HLA genes across microglia/macrophages subsets. The dot size is proportional to the fraction of cells expressing specific genes. The color intensity corresponds to the relative expression of specific genes.

**Figure S4.**
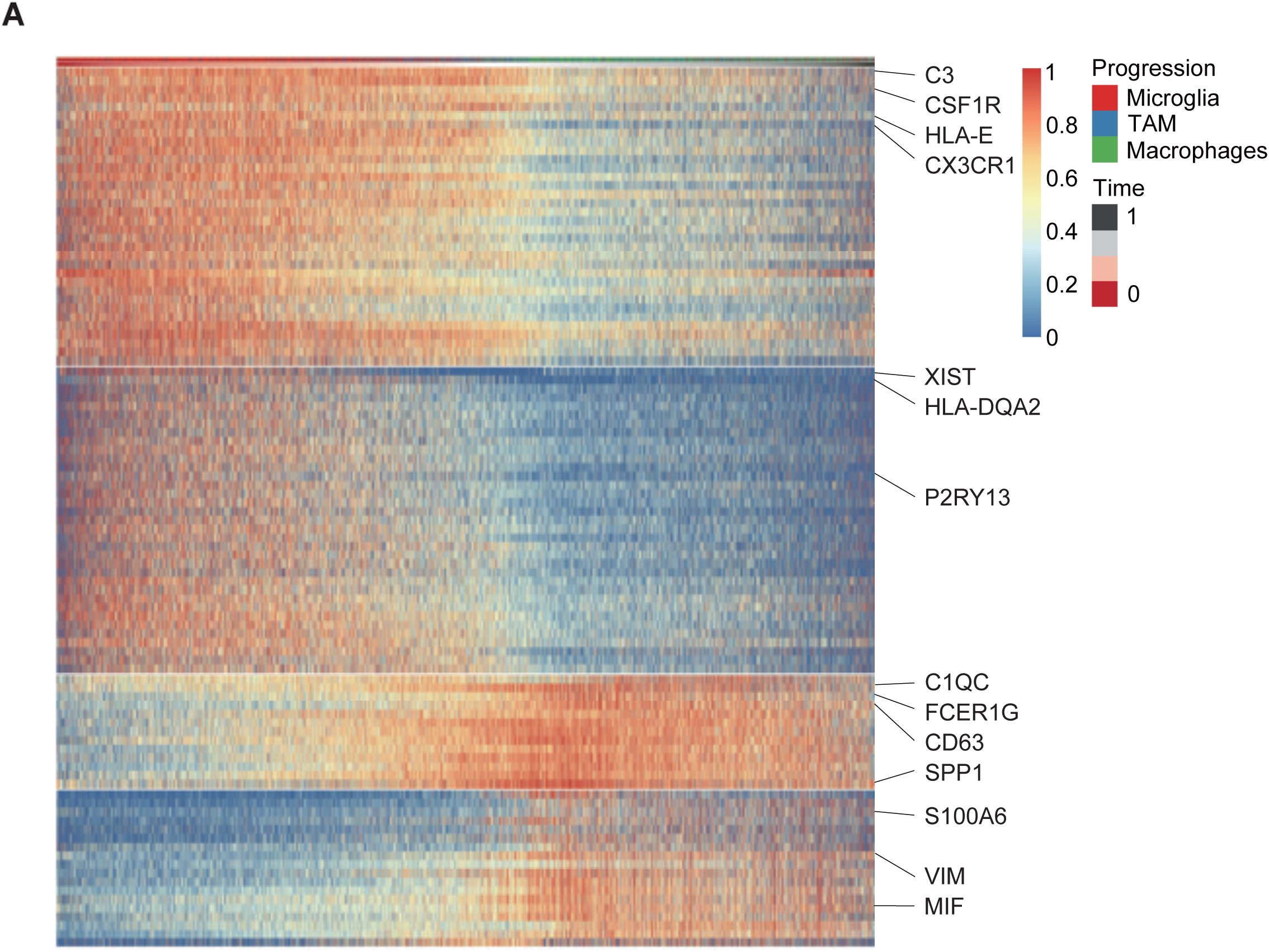
Heatmap showing the differential expression of transcription factors along the pseudo time trajectory of the microglia/macrophage cluster.

**Figure S5.**
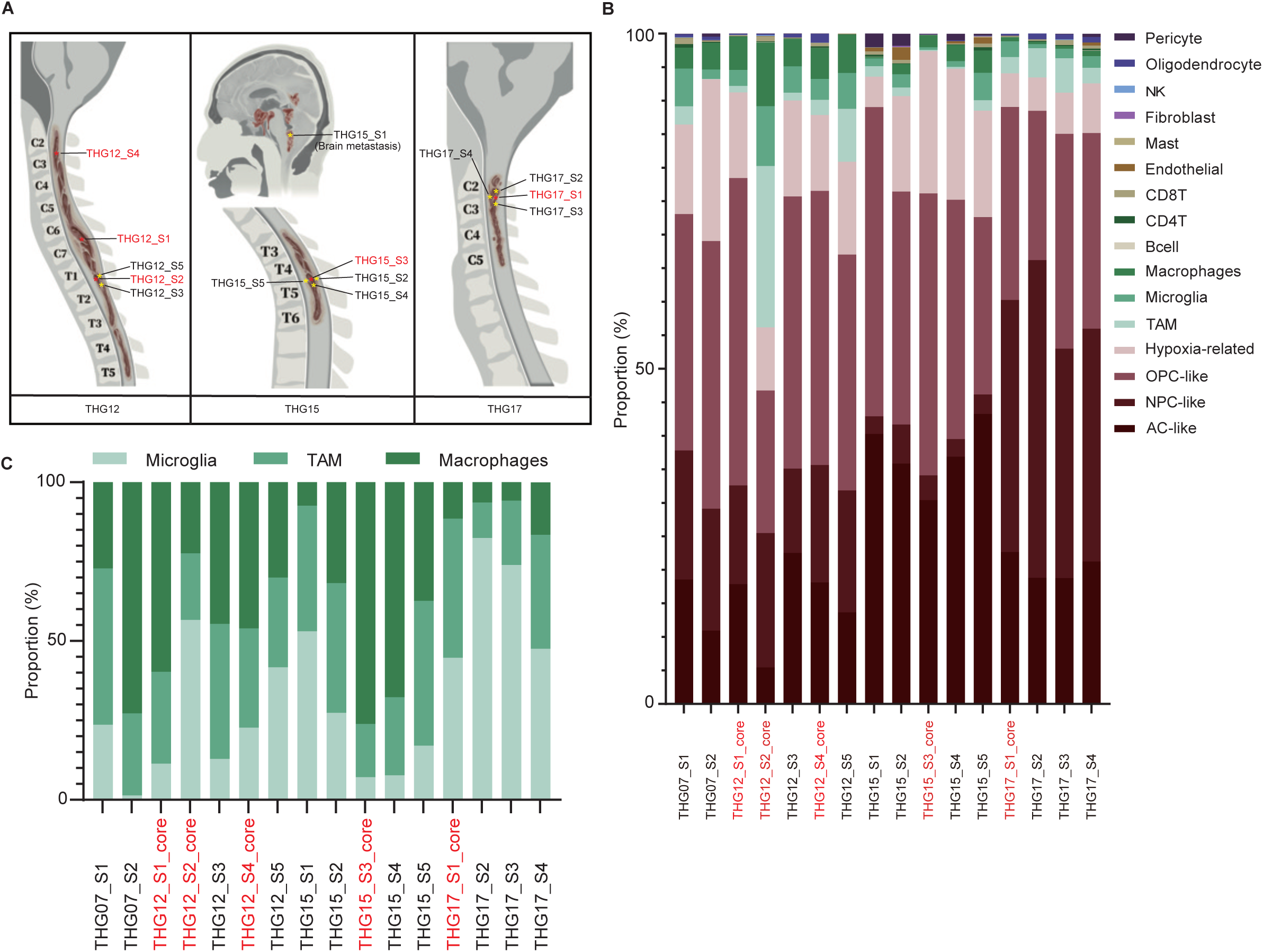
(A) Multiple samples from the same lesion are indicated. The core tumor samples are highlighted in red. (B) The percentages of different subsets of cells in each tumor sample. (C) The percentages of microglia, TAM and macrophages in each tumor sample.

**Figure S6.**
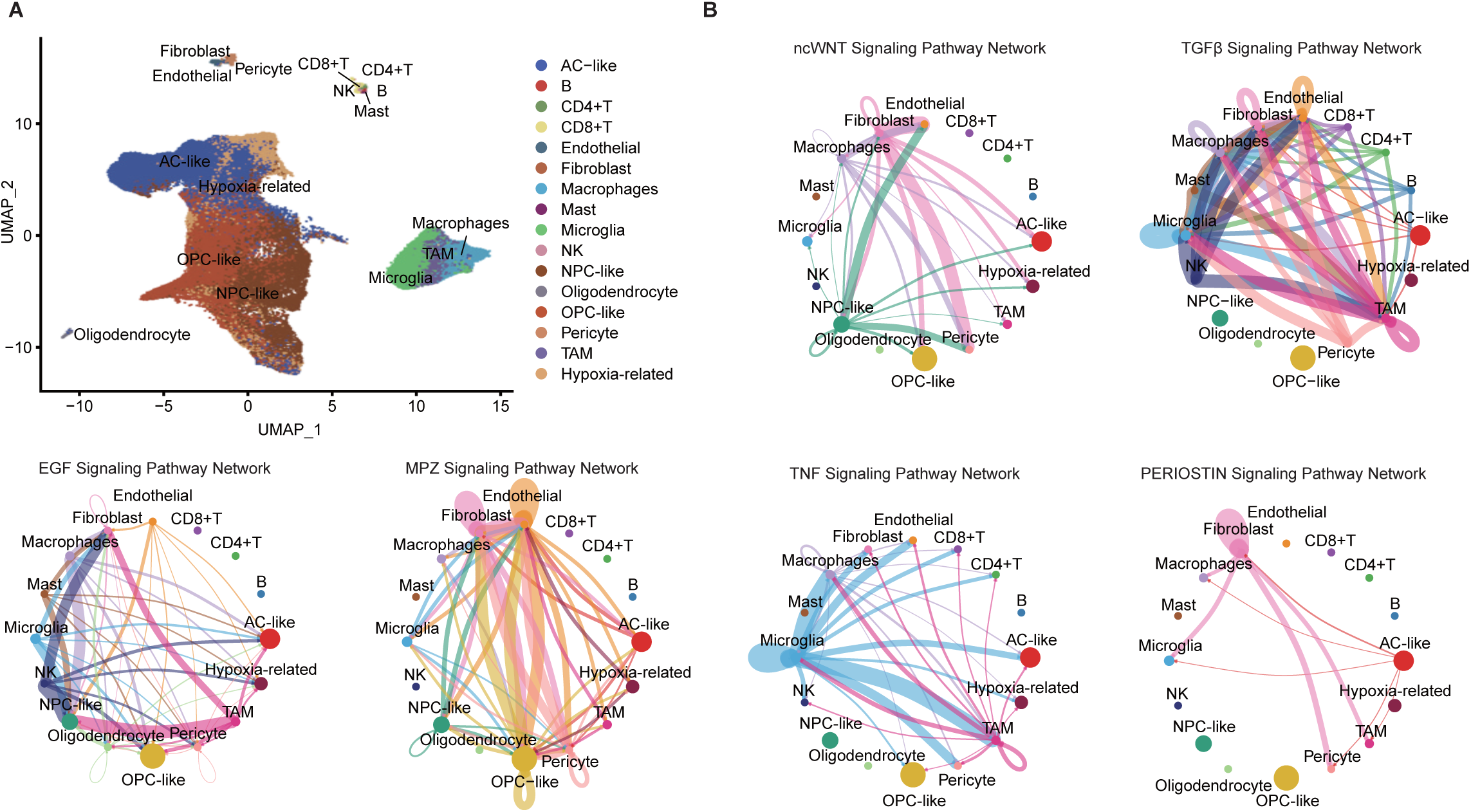
(A) UMAP plot showing all cell subclusters in spinal cord H3K27-altered DMG, colored based on the cell subset. (B) Chord diagrams showing predicted interactions of the TNF, PERIOSTIN, ncWNT, TGFβ, EGF and MPZ signaling pathway networks in spinal cord H3K27-altered DMG.

## Methods

### 1 Patients

To decipher genomic and transcriptional signatures of spinal cord H3K27-altered DMG, 19 fresh tumor samples were collected from 7 consecutive surgical (including biopsy) patients at Beijing Tsinghua Changgung Hospital. We obtained appropriate approval from the institutional review board. Pathologic and molecular diagnoses were confirmed by a consortium of subspecialized neuropathologists. Patient characteristics are summarized in Table 1. The patients’ clinical features were typical of spinal cord H3K27-altered DMG, with the age range of 14 to 34 years and median of 25 years. Four patients had tumor samples obtained from multiple lesions (Figure 1). Matched peripheral blood was also collected from each participant.

To assess the overall survival of patients with H3K27-altered DMG of various tumor locations, we compiled survival data of 38 cases of spinal cord H3K27-altered DMG biopsied or surgically treated at Beijing Tsinghua Changgung Hospital, and reviewed literature data for brainstem (N=79) and thalamus (N=31) DMG (Supplementary information, Table S1). For literature data review, relevant articles were systematically searched within PubMed, EMBASE, and Cochran Library. Only original English articles published by up to 2023 were considered. To avoid erroneous omission, literature involving all comparative studies was searched using the following string of medical subject heading (MeSH) terms: (“glioma”[MeSH Terms] OR “glioma”[All Fields]) AND (H3K27M[All Fields] OR K27M[All Fields]). All identified articles were then systematically and independently assessed by two investigators against the inclusion and exclusion criteria as follow:

Inclusion criteria: 1. DMG with pathology consistent with H3K27M; 2. Tumor located in the thalamus or brainstem; 3. Patients were untreated or treated only per conventional protocols (e.g., surgical resection, radiotherapy, temozolomide, and/or bevacizumab).

Exclusion criteria: 1. DMG with multiple intracranial foci; 2. Patients received non-conventional treatment.

### 2 Patient-derived cell cultures

Patient-derived cell cultures were generated with informed consent in compliance with the Ethics Committee of Beijing Tsinghua Changgung Hospital. Briefly, mechanically minced tissues were processed with 0.1% trypsin and 10 U/mL DNaseI at 37°C for 45 min. The tissues were triturated and passed through a 100-μm cell strainer after complete lysis of red blood cells and several washes. Further details have been described in a previous study [1].

### 3 Sample processing

#### DNA isolation for bulk DNA sequencing

Genomic DNA of peripheral blood and tissue samples of patients with spinal cord H3K27-altered DMG were extracted using the QIAamp DNA Mini Kit (QIAGEN) according to the manufacturer’s specification. The concentrations of DNA were quantified using the Qubit HsDNA Kits (Invitrogen) and the qualities of DNA were evaluated with agarose gel electrophoresis.

#### Tissue collection and dissociation for single cell RNA sequencing

Fresh tumor tissues were collected during surgery and immediately transferred to the laboratory in cold phosphate-buffered saline (PBS) on ice. Tumor tissues were then dissociated mechanically and incubated in digestion buffer which contained 2 mg/ml collagenase IV (Gibco), 10 U/ml DNase I (NEB), and 1 mg/ml papain (Sigma) in PBS. After incubation for about 20–30 min at 37 °C on a thermocycler, 10% fetal bovine serum (Gibco) was used to stop the digestion. Single-cell suspensions dissociated from the tumor were resuspended in 0.04% BSA/PBS and stained with 7-amino-actinomycin D (7-AAD) to sort 7-AAD-negative cells by FACS for single-cell libraries preparation.

#### RNA collection and dissociation for bulk RNA sequencing

Based on the previous report [1], 5×10^5^ THG15 cells were incubated for 48 hours with 1 uM panobinostat, 50 uM tazemetostat, or both in 6-well plates in triplicates. Total RNA was extracted using a MolPure Cell/Tissue Total RNA Kit (Yeasen). Cells were lysed with 350 ul LB buffer after centrifugation and was added to a DNA Removal/RNA Adsorption column to collect the RNA filtrate using BD* solution, W* solution and RNase-free H_2_O.

### 4 Bulk RNA sequencing

#### RNA quantification and qualification, library preparation, clustering and sequencing

RNA integrity was assessed using the RNA Nano 6000 Assay Kit of the Bioanalyzer 2100 system (Agilent Technologies, CA, USA). Total RNA was used as the input material for RNA sample preparation. Briefly, mRNA was purified from the total RNA using poly-T oligo-attached magnetic beads. Fragmentation was carried out using divalent cations under elevated temperature in First Strand Synthesis Reaction Buffer (5X). First strand cDNA was synthesized using random hexamer primer and M-MuLV Reverse Transcriptase, and then using RNaseH to degrade the RNA. Second strand cDNA synthesis was subsequently performed using DNA Polymerase I and dNTP. Remaining overhangs were converted into blunt ends via exonuclease/polymerase activities. After adenylation of the 3’ ends of DNA fragments, Adaptors with hairpin loop structures were ligated to prepare for hybridization. In order to select cDNA fragments of preferentially 370∼420 bp in length, the library fragments were purified with AMPure XP system (Beckman Coulter, Beverly, USA). Then, PCR was performed with Phusion High-Fidelity DNA polymerase, Universal PCR primers and Index (X) Primer. At last, PCR products were purified (AMPure XP system) and library quality was assessed on the Agilent Bioanalyzer 2100 system.

#### Bulk RNA sequencing and analysis

Sequence reads were aligned to the hg38 genome using HISAT2 (version 2.2.1). The coding gene annotation of hg38 was downloaded from ENSEMBL. The reads within exon regions were annotated using featureCounts (version 2.0.3). Transcripts per kilobase million (TPM), calculated from counts of all samples, were used to determine the expression levels using TPMCalculator.

### 5 Whole exome sequencing

A total of 0.6 μg DNA was taken from each sample and sheared into fragments of 180∼280 bp using a Covaris S2 ultrasonicator (Covaris). Agencourt SPRIselect Kit Size Selection was used to purify DNA fragments of the desired size for library preparation. The library was constructed using the NEBNext Ultra DNA Library Prep Kit for Illumina (New England Biolabs) according to the manufacturer’s protocol. Exome regions were captured with Agilent SureSelect Human All Exon V6 (Agilent Technologies) according to the manufacturer’s protocol. The post-hybridization amplification product (2 × 150-bp paired-end reads) was quality-checked and sequenced using Illumina Novaseq PE150 instruments (Illumina). Tumor DNA was sequenced to 200X coverage.

#### Mutation calling with WES Data

Paired reads were mapped to GRCh38 using BWA, followed by BAM sorting, merging of read group BAMs into a single BAM by Samtools, and then duplicate-marking using Picard. We followed the GATK Best Practices (https://software.broadinstitute.org/gatk/best-practices/) to process all BAMs. We used MuTect2 to generate raw Somatic Variant Call Format files (VCFs) and VCFs were further processed by caller-specific filters to tag low quality variants in the FILTER column in VCF. These VCFs were then annotated using Variant Effect Predictor to generate Annotated Somatic VCFs. (https://www.ensembl.org/info/docs/tools/vep/).

#### Copy number alterations

Sequenza (v2.2.0) was used to call copy number alterations (CNA) while considering both ploidy and cellularity [2]. Briefly, we used BAM files from the WES data of each tumor and the paired normal samples as input to calculate the depth ratio, which was normalized based on both GC content bias and the data ratio. We used the default parameter setting to acquire segmented copy numbers and estimate cellularity and ploidy. For each tumor sample, the copy numbers of segments were then divided by ploidy following log2 transformation. After filtering out segments smaller than 500 kb, copy number states were determined for each segment. Copy number gains and losses were defined as at least one copy more and one copy less than the estimated ploidy, respectively. Software and codes are available at https://sequenzatools.bitbucket.io/#/home.

CNVkit (v0.9.2) was also performed with default parameters on the paired tumor-normal WES data [3]. After segmentation, the absolute integer copy number of each segment was estimated using the threshold and clonal methods. Software and codes are available at https://github.com/etal/cnvkit. GISTIC2.0 (Genomic Identification of Significant Targets in Cancer v2.0.23) was used to identify focal gain and loss regions (q < 0.25) [4] Software and codes are available at https://doi.org/www.broadinstitute.org/cancer/pub/GISTIC2.

#### Genomic landscape analysis

To allow for easier investigation of variants and annotations, the VCFs were transformed into project-level tab-delimited Mutation Annotation Format (MAF) files. This was done with a customized tool based upon VCF2MAF. Software and codes are available at https://github.com/mskcc/vcf2maf. We used the MAF files to draw the genomic landscapes of all samples using Maftools (v2.10.0) [5]. Software and codes are available at https://github.com/PoisonAlien/Maftools.

### 6 Single-cell RNA-seq sequencing

#### Single-cell RNA-seq data generation and processing

For all samples, single cells were processed through the 10X Chromium 3’ Single Cell Platform using the Chromium Single Cell 3’ Library, Gel Bead and Chip Kits (10X Genomics, Pleasanton, CA), following the manufacturer’s protocol. Thousands of cells were added to each channel of a chip to be partitioned into Gel Beads in Emulsion (GEMs) in the Chromium instrument, followed by cell lysis and barcoded reverse transcription of RNA in the droplets. Breaking of the emulsion was followed by amplification, fragmentation and addition of adapter and sample index. Sequencing was performed with Illumina (HiSeq 2000).

#### Pre-processing of expression matrices

The raw scRNAseq fastq files were processed using Cell Ranger (v4.0.0) from 10X Genomics Technology and aligned to the GRCh38 reference genome. Bam files and filtered expression matrices were generated using ‘cellranger_count’. All expression matrices were loaded into R 4.0.3 using the ‘Read10X’ function from the Seurat (v4.0.0) [5]. The latter library was used to perform the analysis. Software and codes are available at https://satijalab.org/seurat/.

#### Cell doublet detection and removal

We applied scDblFinder (v1.8.0) in human 10× dataset to identify artifactual libraries generated from two or more cells, where they entered the same microfluidic droplet and were labeled with barcodes. We removed potential doublets for further analysis. Software and codes are available at https://github.com/plger/scDblFinder.

#### Quality control filtering

Pre-processing steps were applied to remove low-quality cells if they met the following criteria: (1) <1000 UMIs (unique molecular identifiers), (2) >9000 or <2000 genes, (3) >25% UMIs derived from the mitochondrial genome, or (4) log10 (genes/UMIs) <0.8. Genes were chosen for downstream analyses if they were detected with more than 5 transcripts by more than 5 cells. Following the quality control procedure, the dataset consisted of 119,967 cells. The filtered gene expression matrix was normalized using Seurat’s NormalizeData function, in which the number of UMIs of each gene was divided by the sum of the total UMIs per cell, multiplied by 10,000, and then transformed to log-scale (In (UMI-per-10000+1)).

#### Selection of highly variable genes (HVGs) and batch effect correction

HVGs were calculated using Seurat function FindVariableFeatures(). For batch effect correction, the corrected fastMNN embeddings were used to perform unsupervised clustering of the cells using graph-based clustering based on SNN-Cliq and PhenoGraph with Seurat functions RunFastMNN. Principal component analysis (PCA) was performed on 30 PCs under the ‘‘mnn’’ mode from the HVGs only. Correction was performed first among the 19 samples. For visualization, the dimensionality of each dataset was further reduced using Uniform Manifold Approximation and Projection (UMAP) with Seurat functions RunUMAP. The PCs used to calculate the embeddings were the same as those used for clustering. Analysis in patient THG15 used the same parameters.

#### Cell type annotation and cluster marker identification

The FindAllMarkers function in Seurat was used to discover markers for each of the identified clusters. Clusters were then classified and annotated on the basis of expression of canonical markers of particular cell types. Method MAST was used to perform differential analysis.

#### UMAP analysis and identification of non-malignant cell types

On analysis of cells in samples from patient THG15, three small clusters were apparent, which were associated with high expression of markers for three non-malignant cell types. We thus identified the cell types based on the respective marker genes. Specifically, microglia/macrophages were identified with expression of CD14, CX3CR1, and CD163. Lymphocytes and mast cells with expression of CD3D, CD3E, and CD3G. One stromal cluster was identified by expression of RGS5, VWF, and ACTA2. On analysis of all sample integrated data, another small cluster was apparent besides the three described above: oligodendrocytes with expression of MBP, PLP1, MAG, and MOG.

#### Subclustering of major cell types

For each major cell type, cells were extracted from the overview integrated dataset first. Next, Lymphocytes and mast cells cluster and stromal clusters were integrated for further subclustering. After integration, genes were scaled to unit variance. Normalizing, batch effect, and clustering were performed as described above. For subclustering of the immune cells cluster, in addition to CD8+ T cells (with markers such as CD3D and CD8) and CD4+ T cells (e.g., CD3D, CD4, IL7R), we defined three small populations through known markers representing mast cells (e.g., MS4A2), B cells (e.g., CD79A) and NK cells (e.g., NCAM1, KLRC1, TRDC). For subclustering of the stromal cluster, pericyte cells (e.g., RGS5, NDUFA4L2), endothelial cells (e.g., PECAM1, VWF) and fibroblast cells (e.g., COL1A1, CST3) were defined through known markers.

#### CNA inference from single-cell data

CNAs were estimated by an integrative Bayesian segmentation approach CopyKAT (v1.0.5) for each sample [6]. TPM gene expression matrix was extracted from the Seurat object. Cells classified to each of the non-malignant cell types above were used as known normal cell to perform Bayesian classification for distinguishing normal cells from malignant cells. Quality control filtering was performed to select the highest quality cells by only including malignant cells with at least five genes in each chromosome to calculate DNA copy numbers. Software and codes are available at https://github.com/navinlabcode/copykat/.

#### Integrated definition of malignant cells

We combined the CNA classification with the UMAP-based classifications, such that the final list of malignant cells included those which were classified as malignant by CopyKAT and were part of the malignant UMAP cluster. We extracted these malignant cells for downstream analyses.

#### Expression programs of intra-tumoral heterogeneity

We used a modified intra-tumoral heterogeneity analyses workflow as a previous publication described [7]. We analyzed tumor cells from each sample separately using NMF to identify programs of expression heterogeneity. NMF was applied to the relative expression values, by transforming all negative values to zero. We performed NMF with the number of factors k ranging from 5-10, and initially defined expression programs as the top 30 genes (by NMF score) for each k. For each sample, we sought robust expression programs by selecting those with an overlap of at least 25 out of 30 genes with a program obtained using a different value of k. To avoid redundancies, from each set of overlapping programs from one sample, we only kept one program, selected based on having the highest overlap with a NMF program identified in another sample.

In order to identify recurrently heterogenous expression programs across samples, we compared expression programs derived from NMF, separately, by hierarchical clustering, using 30 minus the number of overlapping genes as a distance metric. Given the high number of NMF programs, clustering was restricted to programs with at least a minimum overlap of 5 out of 30 genes with a program observed in another sample. Jaccard indices, reflecting the overlap between pairs of signatures, were used for hierarchical clustering of the programs by average linkage. Five major groups of programs were identified by cluster dendrogram in programs from spinal cord H3K27-altered DMG, and six were identified in the comparison with brain H3K27-altered DMG. For each group of programs, a meta-module was then defined as all genes included in at least 17% of the constituent programs. We assessed the functional enrichment of meta-modules using the Metascape webtool (www.metascape.org) [8].

#### Definition of signature scores and assignment of cells to meta-modules

We used cell scores (SC) to evaluate the degree to which individual cells expressed a certain predefined expression gene set using the AddModuleScore() function in Seurat in each cell. Malignant cells were assigned to the highest scoring meta-module, including the four “identity” meta-modules (NPC-like, AC-like, OPC-like, hypoxia-related) but excluding the cell cycle meta-module.

#### Two-dimensional representation of malignant cellular states

The two-dimensional cell-state representation map was created as described by Neftel et al [9]. Briefly, cells were first separated into OPC-like/NPC-like vs. AC-like/Hypoxia-related by the sign of D = max (SCopc,SCnpc) – max(SCac,SCvegf), and D defined the Y-axis of all cells. Next, for OPC-like/NPC-like cells (i.e. D>0), the X-axis value was defined as log2(|SCopc – SCnpc|+1) and for AC-like/Hypoxia-related cells (i.e. D<0), the X-axis was defined as log2(|SCac – SCvegf|).

#### Trajectory inference of malignant cells

We performed scVelo(v0.2.2), CellRank (v1.5.0) and PAGA to infer trajectory in malignant cells [10, 11].

#### scVelo

For malignant cells from all sample integrated data, we used RNV velocity to infer trajectory. CellRanger BAM files were sorted with samtools. Per sample loom files were generated from sorted BAM files using Velocyto. Loom files were merged with loompy (v3.0.6) and subset to contain cells consistent with the Seurat object. Software and codes are available at https://github.com/linnarssonlab/loompy. For scVelo analysis, genes in the merged loom file were filtered for a minimum count of 20 and high variability (dispersion). Count matrices were normalized by counts per cell, and log transformed log (x + 1). Filtering and normalization was applied in the same vein to spliced/unspliced counts. We ran the dynamical model using default parameters to learn the full transcriptional dynamics of splicing kinetics. RNA velocity plots on two-dimensional states coordinates created above were generated with the scvelo.pl.velocity_embedding_stream function. Software and codes are available at https://scvelo.readthedocs.io/.

### Analysis of microglia/macrophage cluster

#### Subcluster of microglia/macrophage

PCA was performed on about 2,000 variable genes in the microglia/macrophages cluster. PC1 genes could be used to distinguish microglia from macrophages. We used the function FindClusters on PC1 to annotate subclusters using known markers of microglia and macrophages.

#### Trajectory inference using SCORPIUS

SCORPIUS (v1.0.8) trajectory inference was performed on the microglia/macrophages cluster. SCORPIUS was run on the log2-normalized counts using the proposed workflow. Software and codes are available at https://github.com/rcannood/SCORPIUS.

#### Gene set variation analysis (GSVA)

Pathway analyses were predominantly performed on the 50 hallmark pathways described in the molecular signature database, exported using the MSigDB database (v7.4). We also assessed the KEGG subset and REACTOME subset of Canonical pathways. To assign pathway activity estimates to individual cells, we applied the GSVA using standard settings, as implemented in the GSVA (v1.30.0) [12]. To assess differential activities of pathways between sub-clusters of cells, we contrasted the activity scores for each cell using Limma (v3.38.3). Differential activities of pathways were calculated for each identified cluster.

#### Pathway activity analysis

PROGENy (v1.51.1) was used to infer pathway activities, coupled to foot-print based enrichment analysis statistic [13]. We used the PROGENy model to predict 14 pathway activities among four tumor meta-modules. Software and codes are available at https://github.com/saezlab/progeny-py.

#### Transcription factor (TF) inference

DoRothEA (v1.0.0) was used to decipher the variation of TF activity among four tumor meta-modules [14]. We loaded the TF regulons and chose “A”, “B”, and “C” high-confidence TF selection in the DoRothEA network. VIPER scores were computed on the DoRothEA regulons, scaled and added as the ‘DoRothEA’ slot on the loom object. To allow for comparison of TF score activities, the mean and standard deviation values of the scaled VIPER scores were computed per group. TFs were ranked according to the variance of their corresponding VIPER scores. The top 5 highly variable scores per tumor meta-module were kept for visualization of their corresponding scores. Software and codes are available at https://saezlab.github.io/dorothea/.

#### Analysis of intercellular communication networks

Ligand-Receptor interactions were analyzed using the CellChat (v1.0.0) based on the database of interactions among ligands, receptors and their cofactors that accurately represent known heteromeric molecular complexes [15]. We analyzed the communication networks among malignant cells and non-malignant cells in the tumor microenvironment. Software and codes are available at https://github.com/sqjin/CellChat.

#### Public brain H3K27-altered DMG scRNA-seq datasets

We reanalyzed the public scRNA-seq data on brain H3K27-altered DMG to dissect variation in the transcriptional landscapes between spinal cord and intracranial DMG [16]. Processed scRNA-seq data were downloaded through the Gene Expression Omnibus (GEO, accession no. GSE102130). We used methods provided in the publication to label cells and isolated tumor cells. Tumor cells from six brain DMG patients were analyzed using intra-tumoral heterogeneity analyses as described above and were compared to our data. Hierarchical clustering of the programs reproduced the four compartments (i.e., Cell cycle, OPC-like, AC-like, and OC-like) defined in the original article with marked genes. Meanwhile, the hypoxia-related and NPC-like meta-modules were lacking in the brain glioma.

### 7 Western blotting assay

To analyze feature gene alterations in drug-exposed tumor cells, 5×10^5^ THG15 cells were incubated for 48 hours with 1 uM Panobinostat and 50 uM tazemetostat in 6-well plates in triplicates. Cell protein was isolated and harvested in RIPA buffer supplemented with protease and phosphatase inhibitors and quantified by BCA assay according to the standard protocols. Equal amounts of protein lysates from each sample were fractionated on 10% Bis-Tris gels. Separated proteins were transferred to an Immobilon^®^-P PVDF Membrane (Millipore) using an eBlot^®^ L1 Fast Wet Transfer System (Genscript). After blockade with 5% skimmed milk dissolved in TBST composed of 1×TBS (Lablead) and 0.1% TWEEN 20 (VWR), protein blots were incubated with primary antibodies against VEGFA (Proteintech), S100A11 (Proteintech), GADD45A (Proteintech), and SHMT2 (Proteintech), followed by horseradish peroxidase-conjugated goat anti-rabbit (EASYBIO) or anti-mouse (EASYBIO) secondary antibodies. Probed blots were detected using SuperSignal^TM^ West Pico PLUS Chemiluminescent Substrate (Thermo Fisher Scientific) and imaged on ImageQuant LAS 4000 (GE). The relative intensities of blots were quantified with densitometric analysis on ImageJ from the National Institutes of Health.

### 8 Immunofluorescence

Frozen 7-uM-thick sections were allowed to dry on Adhesion Microscopeslides (CITOTEST) at room temperature and then were rehydrated in PBS. Sections were blocked in 10% normal goat serum (Thermal Fisher) in PBS + 0.2% Triton X-100 for 1 hour at room temperature. Samples were incubated at 4℃ with the following primary antibodies in PBS + 5% goat serum + 0.2% Triton X-100: VEGFA (Proteintech), GFAP (Proteintech), PDGFRA (Proteintech), and SOX11 (Proteintech). Excess antibody was removed by rinsing in PBS for 5 min for 3 times. Samples were then incubated at room temperature for 1 h with the following secondary fluorescently labeled antibodies: Cy3-conjugated Goat anti-Rabbit IgG (ABclonal) diluted in PBS + 5% goat serum + 0.2% Triton X-100. Excess antibody was removed by rinsing in PBS for 5 min for 3 times. Slides were mounted in antifading mounting medium with DAPI (Solarbio) and imaged with an inverted fluorescence microscope (Ts2R-FL, Nikon) to obtain images.

### 9 Statistics and reproducibility

Plotting and statistical analysis were performed in the R statistical environment (v.4.0.3). The statistical tools, methods, and thresholds for each analysis were explicitly described with the results or detailed in the figure legends or the methods above.

## Notes

### Competing Interest Statement

The authors have declared no competing interest.

## References

[1] Phillips R E, Soshnev A A, Allis C D. Epigenomic Reprogramming as a Driver of Malignant Glioma [J]. Cancer Cell, 2020, 38(5): 647–60.

[2] Louis D N, Perry A, Wesseling P, et al. The 2021 WHO Classification of Tumors of the Central Nervous System: a summary [J]. Neuro Oncol, 2021, 23(8): 1231–51.

[3] Argersinger D P, Rivas S R, Shah A H, et al. New Developments in the Pathogenesis, Therapeutic Targeting, and Treatment of H3K27M-Mutant Diffuse Midline Glioma [J]. Cancers (Basel), 2021, 13(21).

[4] Lowe B R, Maxham L A, Hamey J J, et al. Histone H3 Mutations: An Updated View of Their Role in Chromatin Deregulation and Cancer [J]. Cancers (Basel), 2019, 11(5).

[5] Larson J D, Kasper L H, Paugh B S, et al. Histone H3.3 K27M Accelerates Spontaneous Brainstem Glioma and Drives Restricted Changes in Bivalent Gene Expression [J]. Cancer Cell, 2019, 35(1): 140–55 e7.

[6] Nagaraja S, Vitanza N A, Woo P J, et al. Transcriptional Dependencies in Diffuse Intrinsic Pontine Glioma [J]. Cancer Cell, 2017, 31(5): 635–52 e6.

[7] Jing L, Qian Z, Gao Q, et al. Diffuse midline glioma treated with epigenetic agent-based immunotherapy [J]. Signal Transduct Target Ther, 2023, 8(1): 23.

[8] Cooney T M, Lubanszky E, Prasad R, et al. Diffuse midline glioma: review of epigenetics [J]. J Neurooncol, 2020, 150(1): 27–34.

[9] Price G, Bouras A, Hambardzumyan D, et al. Current knowledge on the immune microenvironment and emerging immunotherapies in diffuse midline glioma [J]. EBioMedicine, 2021, 69: 103453.

[10] Solomon D A, Wood M D, Tihan T, et al. Diffuse Midline Gliomas with Histone H3-K27M Mutation: A Series of 47 Cases Assessing the Spectrum of Morphologic Variation and Associated Genetic Alterations [J]. Brain Pathol, 2016, 26(5): 569–80.

[11] Cooney T, Lane A, Bartels U, et al. Contemporary survival endpoints: an International Diffuse Intrinsic Pontine Glioma Registry study [J]. Neuro Oncol, 2017, 19(9): 1279–80.

[12] Karremann M, Gielen G H, Hoffmann M, et al. Diffuse high-grade gliomas with H3 K27M mutations carry a dismal prognosis independent of tumor location [J]. Neuro Oncol, 2018, 20(1): 123–31.

[13] Zheng L, Gong J, Yu T, et al. Diffuse Midline Gliomas With Histone H3 K27M Mutation in Adults and Children: A Retrospective Series of 164 Cases [J]. Am J Surg Pathol, 2022, 46(6): 863–71.

[14] Wang Z, Wang Z, Zhang C, et al. Genetic and clinical characterization of B7-H3 (CD276) expression and epigenetic regulation in diffuse brain glioma [J]. Cancer Sci, 2018, 109(9): 2697–705.

[15] Mackay A, Burford A, Carvalho D, et al. Integrated Molecular Meta-Analysis of 1,000 Pediatric High-Grade and Diffuse Intrinsic Pontine Glioma [J]. Cancer Cell, 2017, 32(4): 520–37 e5.

[16] Filbin M G, Tirosh I, Hovestadt V, et al. Developmental and oncogenic programs in H3K27M gliomas dissected by single-cell RNA-seq [J]. Science, 2018, 360(6386): 331–5.

[17] Liu I, Jiang L, Samuelsson E R, et al. The landscape of tumor cell states and spatial organization in H3-K27M mutant diffuse midline glioma across age and location [J]. Nat Genet, 2022, 54(12): 1881–94.

[18] Gessi M, Gielen G H, Dreschmann V, et al. High frequency of H3F3A (K27M) mutations characterizes pediatric and adult high-grade gliomas of the spinal cord [J]. Acta Neuropathol, 2015, 130(3): 435–7.

[19] Schreck K C, Ranjan S, Skorupan N, et al. Incidence and clinicopathologic features of H3 K27M mutations in adults with radiographically-determined midline gliomas [J]. J Neurooncol, 2019, 143(1): 87–93.

[20] Vallero S G, Bertero L, Morana G, et al. Pediatric diffuse midline glioma H3K27-altered: A complex clinical and biological landscape behind a neatly defined tumor type [J]. Front Oncol, 2022, 12: 1082062.

[21] Alvi M A, Ida C M, Paolini M A, et al. Spinal cord high-grade infiltrating gliomas in adults: clinico-pathological and molecular evaluation [J]. Mod Pathol, 2019, 32(9): 1236–43.

[22] Wang L, Li Z, Zhang M, et al. H3 K27M-mutant diffuse midline gliomas in different anatomical locations [J]. Hum Pathol, 2018, 78: 89–96.

[23] Qiu T, Chanchotisatien A, Qin Z, et al. Imaging characteristics of adult H3 K27M-mutant gliomas [J]. J Neurosurg, 2020, 133(6): 1662–70.

[24] Wang Y Z, Zhang Y W, Liu W H, et al. Spinal Cord Diffuse Midline Gliomas With H3 K27m-Mutant: Clinicopathological Features and Prognosis [J]. Neurosurgery, 2021, 89(2): 300–7.

[25] Duan Z J, Feng J, Yao K, et al. [Clinicopathological characteristics of H3K27-altered diffuse midline glioma and evaluation of NTRK as its therapeutic target] [J]. Zhonghua Bing Li Xue Za Zhi, 2022, 51(11): 1115–22.

[26] Ishi Y, Takamiya S, Seki T, et al. Prognostic role of H3K27M mutation, histone H3K27 methylation status, and EZH2 expression in diffuse spinal cord gliomas [J]. Brain Tumor Pathol, 2020, 37(3): 81–8.

[27] Zhou C, Zhao H, Yang F, et al. Clinical and Genetic Features of Brainstem Glioma in Adults: A Report of 50 Cases in a Single Center [J]. J Clin Neurol, 2021, 17(2): 220–8.

[28] Aihara K, Mukasa A, Gotoh K, et al. H3F3A K27M mutations in thalamic gliomas from young adult patients [J]. Neuro Oncol, 2014, 16(1): 140–6.

[29] Bruzek A K, Ravi K, Muruganand A, et al. Electronic DNA Analysis of CSF Cell-free Tumor DNA to Quantify Multi-gene Molecular Response in Pediatric High-grade Glioma [J]. Clin Cancer Res, 2020, 26(23): 6266–76.

[30] Khadka P, Reitman Z J, Lu S, et al. PPM1D mutations are oncogenic drivers of de novo diffuse midline glioma formation [J]. Nat Commun, 2022, 13(1): 604.

[31] Enomoto T, Aoki M, Hamasaki M, et al. Midline Glioma in Adults: Clinicopathological, Genetic, and Epigenetic Analysis [J]. Neurol Med Chir (Tokyo), 2020, 60(3): 136–46.

[32] Schulte J D, Buerki R A, Lapointe S, et al. Clinical, radiologic, and genetic characteristics of histone H3 K27M-mutant diffuse midline gliomas in adults [J]. Neurooncol Adv, 2020, 2(1): vdaa142.

[33] Neftel C, Laffy J, Filbin M G, et al. An Integrative Model of Cellular States, Plasticity, and Genetics for Glioblastoma [J]. Cell, 2019, 178(4): 835–49 e21.

[34] Kinker G S, Greenwald A C, Tal R, et al. Pan-cancer single-cell RNA-seq identifies recurring programs of cellular heterogeneity [J]. Nat Genet, 2020, 52(11): 1208–18.

[35] Pachocki C J, Hol E M. Current perspectives on diffuse midline glioma and a different role for the immune microenvironment compared to glioblastoma [J]. J Neuroinflammation, 2022, 19(1): 276.

[36] Venteicher A S, Tirosh I, Hebert C, et al. Decoupling genetics, lineages, and microenvironment in IDH-mutant gliomas by single-cell RNA-seq [J]. Science, 2017, 355(6332).

[37] Zhou W, Ke S Q, Huang Z, et al. Periostin secreted by glioblastoma stem cells recruits M2 tumour-associated macrophages and promotes malignant growth [J]. Nat Cell Biol, 2015, 17(2): 170–82.

[38] Guo X, Xue H, Shao Q, et al. Hypoxia promotes glioma-associated macrophage infiltration via periostin and subsequent M2 polarization by upregulating TGF-beta and M-CSFR [J]. Oncotarget, 2016, 7(49): 80521–42.

[39] Yan S, Han X, Xue H, et al. Let-7f Inhibits Glioma Cell Proliferation, Migration, and Invasion by Targeting Periostin [J]. J Cell Biochem, 2015, 116(8): 1680–92.

[40] Wierzbicki K, Ravi K, Franson A, et al. Targeting and Therapeutic Monitoring of H3K27M-Mutant Glioma [J]. Curr Oncol Rep, 2020, 22(2): 19.

[41] Mohammad F, Weissmann S, Leblanc B, et al. EZH2 is a potential therapeutic target for H3K27M-mutant pediatric gliomas [J]. Nat Med, 2017, 23(4): 483–92.

[42] Lin G L, Wilson K M, Ceribelli M, et al. Therapeutic strategies for diffuse midline glioma from high-throughput combination drug screening [J]. Sci Transl Med, 2019, 11(519).

[43] Neth B J, Balakrishnan S N, Carabenciov I D, et al. Panobinostat in adults with H3 K27M-mutant diffuse midline glioma: a single-center experience [J]. J Neurooncol, 2022, 157(1): 91–100.

[44] Peters K, Pratt D, Koschmann C, et al. Prolonged survival in a patient with a cervical spine H3K27M-mutant diffuse midline glioma [J]. BMJ Case Rep, 2019, 12(10).

[45] Vitanza N A, Biery M C, Myers C, et al. Optimal therapeutic targeting by HDAC inhibition in biopsy-derived treatment-naive diffuse midline glioma models [J]. Neuro Oncol, 2021, 23(3): 376–86.

[46] Kim D, Fiske B P, Birsoy K, et al. SHMT2 drives glioma cell survival in ischaemia but imposes a dependence on glycine clearance [J]. Nature, 2015, 520(7547): 363–7.

[47] Wang H H, Chang T Y, Lin W C, et al. GADD45A plays a protective role against temozolomide treatment in glioblastoma cells [J]. Sci Rep, 2017, 7(1): 8814.

[48] Wang H, Yin M, Ye L, et al. S100A11 Promotes Glioma Cell Proliferation and Predicts Grade-Correlated Unfavorable Prognosis [J]. Technol Cancer Res Treat, 2021, 20: 15330338211011961.

[49] Chen L, Xie X, Wang T, et al. ARL13B promotes angiogenesis and glioma growth by activating VEGFA-VEGFR2 signaling [J]. Neuro Oncol, 2023, 25(5): 871–85.

[50] Gu Q, Huang Y, Zhang H, et al. Case Report: Five Adult Cases of H3K27-Altered Diffuse Midline Glioma in the Spinal Cord [J]. Front Oncol, 2021, 11: 701113.

[51] Dono A, Takayasu T, Ballester L Y, et al. Adult diffuse midline gliomas: Clinical, radiological, and genetic characteristics [J]. J Clin Neurosci, 2020, 82(Pt A): 1–8.

[52] Yao J, Wang L, Ge H, et al. Diffuse midline glioma with H3 K27M mutation of the spinal cord: A series of 33 cases [J]. Neuropathology, 2021, 41(3): 183–90.

[53] Arunachalam S, Szlachta K, Brady S W, et al. Convergent evolution and multi-wave clonal invasion in H3 K27-altered diffuse midline gliomas treated with a PDGFR inhibitor [J]. Acta Neuropathol Commun, 2022, 10(1): 80.

[54] Findlay I J, De Iuliis G N, Duchatel R J, et al. Pharmaco-proteogenomic profiling of pediatric diffuse midline glioma to inform future treatment strategies [J]. Oncogene, 2022, 41(4): 461–75.

[55] Reardon D A, Brandes A A, Omuro A, et al. Effect of Nivolumab vs Bevacizumab in Patients With Recurrent Glioblastoma: The CheckMate 143 Phase 3 Randomized Clinical Trial [J]. JAMA Oncol, 2020, 6(7): 1003–10.

[56] Lim M, Weller M, Idbaih A, et al. Phase III trial of chemoradiotherapy with temozolomide plus nivolumab or placebo for newly diagnosed glioblastoma with methylated MGMT promoter [J]. Neuro Oncol, 2022, 24(11): 1935–49.

[57] Omuro A, Brandes A A, Carpentier A F, et al. Radiotherapy combined with nivolumab or temozolomide for newly diagnosed glioblastoma with unmethylated MGMT promoter: An international randomized phase III trial [J]. Neuro Oncol, 2023, 25(1): 123–34.

[58] Kline C, Liu S J, Duriseti S, et al. Reirradiation and PD-1 inhibition with nivolumab for the treatment of recurrent diffuse intrinsic pontine glioma: a single-institution experience [J]. J Neurooncol, 2018, 140(3): 629–38.

[59] Bartanusz V, Jezova D, Alajajian B, et al. The blood-spinal cord barrier: morphology and clinical implications [J]. Ann Neurol, 2011, 70(2): 194–206.

[60] Sweeney M D, Zhao Z, Montagne A, et al. Blood-Brain Barrier: From Physiology to Disease and Back [J]. Physiol Rev, 2019, 99(1): 21–78.

[61] Muz B, de la Puente P, Azab F, et al. The role of hypoxia in cancer progression, angiogenesis, metastasis, and resistance to therapy [J]. Hypoxia (Auckl), 2015, 3: 83–92.

[62] Chen Z, Han F, Du Y, et al. Hypoxic microenvironment in cancer: molecular mechanisms and therapeutic interventions [J]. Signal Transduct Target Ther, 2023, 8(1): 70.

[63] Goodwin M L, Pennington Z, Westbroek E M, et al. Lactate and cancer: a “lactatic” perspective on spinal tumor metabolism (part 1) [J]. Ann Transl Med, 2019, 7(10): 220.

[64] Yin X, Duan H, Yi Z, et al. Incidence, Prognostic Factors and Survival for Hemangioblastoma of the Central Nervous System: Analysis Based on the Surveillance, Epidemiology, and End Results Database [J]. Front Oncol, 2020, 10: 570103.

[65] Chai R C, Yan H, An S Y, et al. Genomic profiling and prognostic factors of H3 K27M-mutant spinal cord diffuse glioma [J]. Brain Pathol, 2023, 33(4): e13153.

[66] Couturier C P, Ayyadhury S, Le P U, et al. Single-cell RNA-seq reveals that glioblastoma recapitulates a normal neurodevelopmental hierarchy [J]. Nat Commun, 2020, 11(1): 3406.

[67] Chanoch-Myers R, Wider A, Suva M L, et al. Elucidating the diversity of malignant mesenchymal states in glioblastoma by integrative analysis [J]. Genome Med, 2022, 14(1): 106.

[68] Wang L, Babikir H, Muller S, et al. The Phenotypes of Proliferating Glioblastoma Cells Reside on a Single Axis of Variation [J]. Cancer Discov, 2019, 9(12): 1708–19.

[69] Wang L, Jung J, Babikir H, et al. A single-cell atlas of glioblastoma evolution under therapy reveals cell-intrinsic and cell-extrinsic therapeutic targets [J]. Nat Cancer, 2022, 3(12): 1534–52.

[70] Sottoriva A, Spiteri I, Piccirillo S G, et al. Intratumor heterogeneity in human glioblastoma reflects cancer evolutionary dynamics [J]. Proc Natl Acad Sci U S A, 2013, 110(10): 4009–14.

[71] Mathur R, Wang Q, Schupp P G, et al. Glioblastoma evolution and heterogeneity from a 3D whole-tumor perspective [J]. Cell, 2024, 187(2): 446–63 e16.

[72] Miklja Z, Pasternak A, Stallard S, et al. Molecular profiling and targeted therapy in pediatric gliomas: review and consensus recommendations [J]. Neuro Oncol, 2019, 21(8): 968–80.

[73] Liu X, Zhao J, Dong P, et al. TRIM6 silencing for inhibiting growth and angiogenesis of gliomas by regulating VEGFA [J]. J Chem Neuroanat, 2023, 132: 102291.

[74] Hummel T R, Salloum R, Drissi R, et al. A pilot study of bevacizumab-based therapy in patients with newly diagnosed high-grade gliomas and diffuse intrinsic pontine gliomas [J]. J Neurooncol, 2016, 127(1): 53–61.

[75] Evans M, Gill R, Bull K S. Does a Bevacizumab-based regime have a role in the treatment of children with diffuse intrinsic pontine glioma? A systematic review [J]. Neurooncol Adv, 2022, 4(1): vdac100.

[76] Yabuno S, Kawauchi S, Umakoshi M, et al. Spinal Cord Diffuse Midline Glioma, H3K27M-mutant Effectively Treated with Bevacizumab: A Report of Two Cases [J]. NMC Case Rep J, 2021, 8(1): 505–11.

[77] Kumar A, Rashid S, Singh S, et al. Spinal Cord Diffuse Midline Glioma in a 4-Year-Old Boy [J]. Child Neurol Open, 2019, 6: 2329048X19842451.

[78] Su J M, Murray J C, McNall-Knapp R Y, et al. A phase 2 study of valproic acid and radiation, followed by maintenance valproic acid and bevacizumab in children with newly diagnosed diffuse intrinsic pontine glioma or high-grade glioma [J]. Pediatr Blood Cancer, 2020, 67(6): e28283.

[79] Capdevielle C, Desplat A, Charpentier J, et al. HDAC inhibition induces expression of scaffolding proteins critical for tumor progression in pediatric glioma: focus on EBP50 and IRSp53 [J]. Neuro Oncol, 2020, 22(4): 550–62.

[80] Xu K, Ramesh K, Huang V, et al. Final Report on Clinical Outcomes and Tumor Recurrence Patterns of a Pilot Study Assessing Efficacy of Belinostat (PXD-101) with Chemoradiation for Newly Diagnosed Glioblastoma [J]. Tomography, 2022, 8(2): 688–700.

[81] Su J M, Kilburn L B, Mansur D B, et al. Phase I/II trial of vorinostat and radiation and maintenance vorinostat in children with diffuse intrinsic pontine glioma: A Children’s Oncology Group report [J]. Neuro Oncol, 2022, 24(4): 655–64.

## References

1 Jing, L. et al. Diffuse midline glioma treated with epigenetic agent-based immunotherapy. Signal Transduct Target Ther 8, 23, doi:10.1038/s41392-022-01274-7 (2023).

## References

[1] Jing L, Qian Z, Gao Q, et al. Diffuse midline glioma treated with epigenetic agent-based immunotherapy [J]. Signal Transduct Target Ther, 2023, 8(1): 23.

[2] Jin S, Guerrero-Juarez C F, Zhang L, et al. Inference and analysis of cell-cell communication using CellChat [J]. Nat Commun, 2021, 12(1): 1088.

[3] Gao R, Bai S, Henderson Y C, et al. Delineating copy number and clonal substructure in human tumors from single-cell transcriptomes [J]. Nat Biotechnol, 2021, 39(5): 599–608.

[4] Kinker G S, Greenwald A C, Tal R, et al. Pan-cancer single-cell RNA-seq identifies recurring programs of cellular heterogeneity [J]. Nat Genet, 2020, 52(11): 1208–18.

[5] Holland C H, Tanevski J, Perales-Paton J, et al. Robustness and applicability of transcription factor and pathway analysis tools on single-cell RNA-seq data [J]. Genome Biol, 2020, 21(1): 36.

[6] Bergen V, Lange M, Peidli S, et al. Generalizing RNA velocity to transient cell states through dynamical modeling [J]. Nat Biotechnol, 2020, 38(12): 1408–14.

[7] Zhou Y, Zhou B, Pache L, et al. Metascape provides a biologist-oriented resource for the analysis of systems-level datasets [J]. Nat Commun, 2019, 10(1): 1523.

[8] Wolf F A, Hamey F K, Plass M, et al. PAGA: graph abstraction reconciles clustering with trajectory inference through a topology preserving map of single cells [J]. Genome Biol, 2019, 20(1): 59.

[9] Neftel C, Laffy J, Filbin M G, et al. An Integrative Model of Cellular States, Plasticity, and Genetics for Glioblastoma [J]. Cell, 2019, 178(4): 835–49 e21.

[10] Mayakonda A, Lin D C, Assenov Y, et al. Maftools: efficient and comprehensive analysis of somatic variants in cancer [J]. Genome Res, 2018, 28(11): 1747–56.

[11] Filbin M G, Tirosh I, Hovestadt V, et al. Developmental and oncogenic programs in H3K27M gliomas dissected by single-cell RNA-seq [J]. Science, 2018, 360(6386): 331–5.

[12] Talevich E, Shain A H, Botton T, et al. CNVkit: Genome-Wide Copy Number Detection and Visualization from Targeted DNA Sequencing [J]. PLoS Comput Biol, 2016, 12(4): e1004873.

[13] Satija R, Farrell J A, Gennert D, et al. Spatial reconstruction of single-cell gene expression data [J]. Nat Biotechnol, 2015, 33(5): 495–502.

[14] Favero F, Joshi T, Marquard A M, et al. Sequenza: allele-specific copy number and mutation profiles from tumor sequencing data [J]. Ann Oncol, 2015, 26(1): 64–70.

[15] Hanzelmann S, Castelo R, Guinney J. GSVA: gene set variation analysis for microarray and RNA-seq data [J]. BMC Bioinformatics, 2013, 14: 7.

[16] Mermel C H, Schumacher S E, Hill B, et al. GISTIC2.0 facilitates sensitive and confident localization of the targets of focal somatic copy-number alteration in human cancers [J]. Genome Biol, 2011, 12(4): R41.

